# ConfDock: Atom-specific Uncertainty Quantification for Molecular Docking via Conformal Prediction

**DOI:** 10.64898/2026.06.29.735353

**Authors:** Haochang Hao, Nour Elhendawy, Yihang Wang, Lu Cheng

## Abstract

Molecular docking is widely used in structure-based drug discovery, yet most approaches provide point estimates without rigorous uncertainty quantification. This limitation makes it difficult to assess when a predicted pose should be trusted, especially when docking methods are applied to diverse protein–ligand systems. We present ConfDock, a conformal prediction (CP) framework for constructing atom-specific prediction intervals for ligand docking poses. ConfDock combines graph neural network (GNN) based quantile estimation with split conformal calibration, producing intervals that adapt to local protein–ligand environments while retaining distribution-free finite-sample coverage guarantees. We evaluate ConfDock on 238 protein–ligand complexes across four docking methods representing distinct computational paradigms. The proposed approach yields substantially narrower prediction intervals compared to standard split CP (57.2% average reduction in mean interval width, up to 74.5%) while maintaining target coverage across all evaluated settings. Ablation analysis indicates that the GNN captures the dominant structure-dependent variability in uncertainty, whereas the conformal calibration step provides a bounded adjustment to ensure coverage guarantees. These results demonstrate that combining learned, structure-aware quantile estimation with conformal calibration enables rigorous uncertainty quantification for molecular docking at atom-level resolution.

## Introduction

The identification of drug candidates increasingly relies on computational predictions of how small molecules bind to protein targets. This process is also known as molecular docking [1, 2]. Modern docking methods, ranging from physics-based methods like AutoDock Vina [3] and commercial docking suites [4], to geometric deep-learning approaches [5, 6], diffusion-based generative models [7, 8], and end-to-end structure prediction systems [9–11], have achieved remarkable accuracy, with leading approaches placing ligands within 2 Å of their experimentally determined positions for a substantial fraction of targets. Yet a fundamental question remains unanswered: *when a docking algorithm predicts a binding pose, how much should we trust that prediction?* In a modern drug discovery pipeline, docking serves as one of the initial filter before resource-intensive molecular dynamics (MD) simulations and in vitro assays. Because most current docking methods are uncalibrated and lack formal statistical guarantees. This can create a downstream bottleneck: researchers might expend substantial computational or experimental resources to evaluate unreliable poses or risk deprioritizing promising ligands whose binding modes are incorrectly predicted.

Nevertheless, most docking algorithms produce point predictions without rigorous measures of uncertainty. While some methods do provide confidence estimates—DiffDock’s learned confidence model [7], GNINA’s CNNscore [12], and AlphaFold3’s pLDDT/ipTM metrics [10]—these scores are architecture-specific, uncalibrated, and lack formal statistical guarantees. Recent studies have further highlighted this issue: Garcia et al. [13] showed that structure prediction confidence scores have limited ability to predict experimental design success, and Wan et al. [14] demonstrated that Boltz-2 can exhibit overconfidence in its structure and binding affinity predictions. Bayesian approaches [15, 16], deep ensemble methods [17], MC dropout [18], and pose ensemble methods [19] offer alternative uncertainty frameworks but impose distributional assumptions or lack formal coverage guarantees. None of these methods can answer the question: “with what probability does this atom’s predicted coordinate fall within a given interval?”

Conformal prediction (CP) [20–23] offers a principled alternative: distribution-free, finite-sample coverage guarantees that require only the mild assumption of data exchangeability (i.e., the joint distribution of the calibration and test data is invariant to permutation). However, standard CP and its variants [24–33] face fundamental limitations in molecular docking. In particular, atom-level errors are heterogeneous and depend on each atom’s local structural environment, which that cannot be adequately represented by the single-valued measures of sample difficulty used by these methods. Although recent work has demonstrated the promise of combining CP with graph neural networks (GNNs) for node/edge-level tasks [34–37] and scalar molecular properties [38–40], these efforts do not address structured spatial predictions where uncertainty varies across atoms within a single complex. This leaves a critical gap: standard CP methods provide rigorous coverage guarantees but yield uninformative, uniform intervals, while learned confidence scores can adapt to local structure but lack formal statistical validity.

Here we introduce ConfDock, a framework that bridges this gap by integrating GNNs with CP for atom-specific uncertainty quantification (UQ) in molecular docking. Our key insight is to use GNNs not as predictors of binding poses, but as learners of conditional quantile functions of docking errors from the structural context of protein-ligand interaction graphs. Specifically, we train a graph attention network [41, 42] with quantile regression loss to predict asymmetric bounds for each atom, then apply conformal calibration to provide finite-sample coverage guarantees. The resulting intervals are simultaneously adaptive (varying across atoms based on local structural context), asymmetric (allowing unequal uncertainty on either side of the predicted coordinate), and statistically valid (carrying distribution-free marginal coverage guarantees).

We evaluate ConfDock on 238 protein-ligand systems across 14 target proteins, drawn from the Open Forcefield protein-ligand binding benchmark [43] (with structures originally from PDB [44]). We consider four representative docking methods spanning distinct computational paradigms: physics-based scoring with AutoDock Vina [3], GNN-augmented refinement with MedusaGraph [45], generative diffusion with DiffDock [7], and end-to-end structure prediction with Protenix [46]. We report several key findings. First, CQR-GNN produces prediction intervals 57.2% narrower than standard split conformal prediction while maintaining valid coverage (≥89%) across all 16 configurations, demonstrating that learning atom-specific uncertainty through molecular graphs yields dramatically more informative intervals. Second, atom-specific intervals reveal a reliability hierarchy across the four docking methods on our benchmark: AutoDock Vina produces the narrowest intervals (4.58 Å mean), followed by Protenix (7.50 Å), DiffDock (7.91 Å), and MedusaGraph (9.81 Å). This hierarchy reflects the error distributions of the evaluated systems rather than a general ranking of docking-method performance, and cannot be obtained from traditional uniform-interval approaches. These results establish ConfDock as the first rigorous, atom-level reliability assessment framework for molecular docking, opening the door to confidence-aware drug discovery pipelines where experimental resources are allocated based on quantified prediction reliability.

## Methods

ConfDock constructs atom-specific prediction intervals for any docking method’s output, with distribution-free, finite-sample coverage guarantees. Given predicted ligand coordinates, the framework (i) represents the protein-ligand complex as an attributed molecular graph, (ii) uses a GNN [47, 48] to learn asymmetric conditional-quantile bounds on each atom’s prediction error, and (iii) applies a conformal calibration step that converts these bounds into intervals with provable marginal coverage. Because the GNN operates only on the predicted coordinates and the resulting graph rather than on the docking algorithm itself, ConfDock is agnostic to the underlying docking method.

The remainder of this section first reviews the necessary background on CP. We then introduce the design rationale of ConfDock, followed by the CQR-GNN architecture and training objective. Finally, we describe the conformal calibration step that maps the GNN’s raw quantile outputs into intervals with formal coverage guarantees.

### Conformal prediction background

We set up the CP framework that ConfDock builds on. Let *X*_*i*_ denote the input features of the *i*-th example (in our setting, the protein-ligand graph together with the index of the target atom), and *Y*_*i*_ denote the corresponding ground-truth scalar target (in our setting, the signed prediction error of that atom along a chosen axis). Given calibration examples 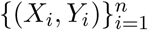 and a new test point (*X*_*n*+1_, *Y*_*n*+1_), CP [20, 21, 23, 49] computes a *nonconformity score s*_*i*_ = *S*(*X*_*i*_, *Y*_*i*_) measuring how poorly the model’s prediction matches the true value. For regression, a common choice is 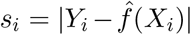, where 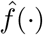 is a pretrained regression model that maps input features *X*_*i*_ to a point prediction of *Y*_*i*_. The prediction interval is:

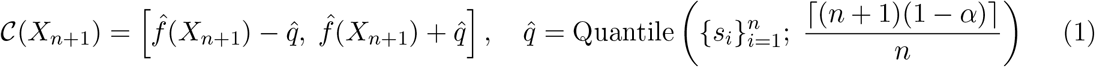

Under the sole assumption that calibration and test data are exchangeable, CP guarantees ℙ (*Y*_*n*+1_ ∈ C(*X*_*n*+1_)) ≥ 1 − *α* for any target miscoverage rate *α* and any data distribution.

### High-level design

The design of ConfDock is motivated by three observations. First, uncertainty in docking predictions is fundamentally determined by local molecular structure, such as binding pocket geometry, protein-ligand contacts, and the local chemical and steric environment of each atom. This information can be naturally encoded in a molecular graph, motivating a GNN backbone. Recent geometric GNNs [50– 53] have demonstrated the effectiveness of learning from 3D molecular structure. Second, docking error distributions are typically asymmetric and heteroscedastic, motivating conformalized quantile regression (CQR) [26] with separate upper and lower quantile bounds. Third, coverage guarantees require a distribution-free calibration step; CP provides exactly this, serving as a “safety net” that adapts its role automatically.

These observations lead to three key design decisions:

#### System-level data splitting

All poses from the same protein-ligand system are assigned to the same partition. Drug-discovery deployments encounter unseen protein targets rather than new poses of already-seen targets, so the realistic test of a UQ method is whether it generalizes across systems.

#### Multi-component training objective

The pinball loss alone has several failure modes in finite-sample training: realized coverage can drift away from the target, the learned embedding need not reflect error structure, and large interval violations contribute the same gradient as small ones once the bound is missed. We therefore augment the pinball term with a coverage-aware width regularization (anchors empirical coverage to the target), a contrastive loss (shapes the embedding by error structure), and a heteroscedastic loss (provides magnitude-aware feedback on bound violations); each component addresses a specific deficiency of the bare pinball objective.

#### Global and local conformal calibration

Atom-level interval widths can vary by an order of magnitude within a single complex (e.g., narrow for buried hydrophobic-core atoms, wide for solvent-exposed flexible atoms). A single global adjustment *η* applied uniformly is then either too lax for narrow-width atoms or too conservative for wide ones. We therefore additionally compute bin-wise adjustments *η*_*b*_ on calibration atoms grouped by predicted width, recovering tighter local guarantees where the heteroscedasticity is pronounced; the better-performing strategy (global vs. bin-wise) is selected per configuration.

#### Pipeline overview

Figure 1 illustrates the three-step ConfDock pipeline. Given predicted ligand coordinates from any docking method, each protein-ligand complex is first represented as an attributed molecular graph with 82-dimensional SE(3)-invariant node features [54] and edges connecting atoms within a 6.0 Å cutoff (Step 1). A graph attention network then propagates information across the molecular graph, learning from protein-ligand structural context which ligand atoms are harder to predict. The edge weights in the figure reflect learned attention importance (Step 2). The GNN outputs a separate lower and upper interval bound for each ligand atom, allowing the interval to be asymmetric around the predicted coordinate. A conformal adjustment, either a single global value or a bin-wise correction stratified by predicted width and calibrated on a held-out set of unseen systems, is then added symmetrically to both sides to provide distribution-free coverage guarantees (Step 3). Because the framework is agnostic to the underlying docking method, it enables direct reliability comparison across different computational paradigms.

**Figure 1.**
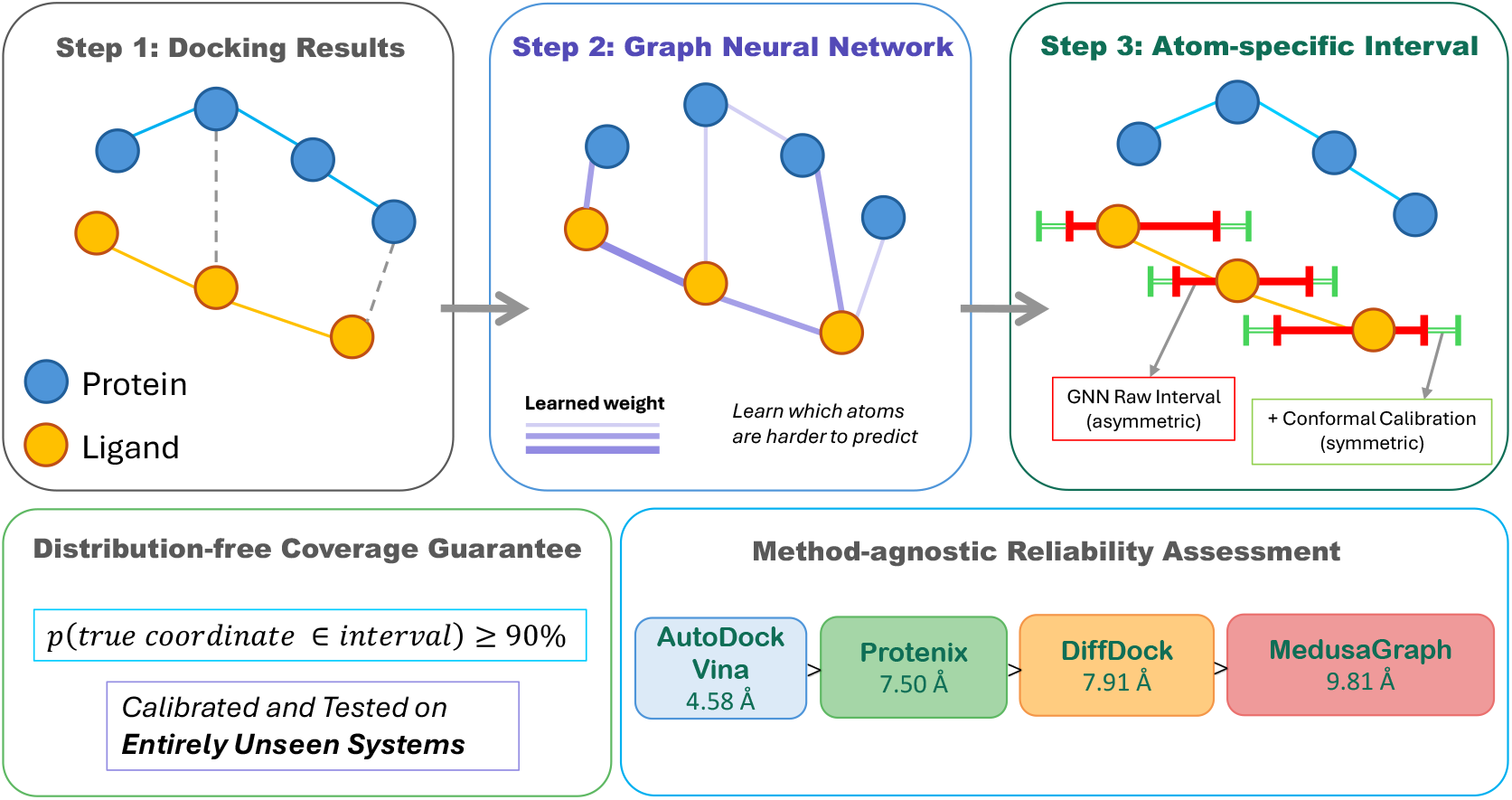
Overview of the ConfDock framework. Docking results are converted into molecular graphs (Step 1), processed by a GNN that learns atom-level error patterns (Step 2), and transformed into asymmetric prediction intervals with conformal coverage guarantees for each ligand atom (Step 3). The framework applies to any docking method, enabling quantitative reliability assessment across computational paradigms.

### CQR-GNN architecture

For each ligand atom *v* ∈ V_*L*_, the docking method produces a predicted coordinate *ŷ*_*v*_ along a specified axis, while 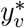 denotes the corresponding ground-truth coordinate measured from the experimentally determined crystal structure. The quantity of interest is the signed prediction error 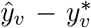: CQR-GNN learns asymmetric bounds 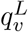 and 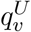 on this error so that the interval 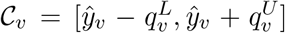 contains the true coordinate with probability at least 1 − *α*, i.e., 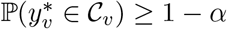.

CQR-GNN employs an enhanced GATv2-based architecture:

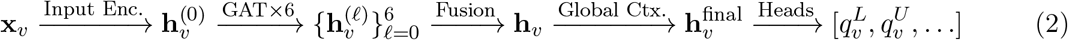

Starting from the 82-dimensional atom feature vector **x**_*v*_, the input encoder produces an initial node embedding 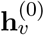 via a two-layer MLP (multi-layer perceptron) with layer normalization, GELU activation [55], and input clamping to handle extreme feature values.

In a protein-ligand graph each ligand atom is connected to many protein atoms within the cutoff distance, but only a subset of those contacts carry information about prediction uncertainty. We therefore choose a graph-attention backbone [41, 42] that learns to down-weight irrelevant neighbors rather than averaging over them uniformly, as plain message-passing or GCN layers would: *L* = 6 enhanced graph-attention blocks successively update the embedding to 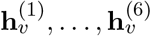, each block combining GATv2 multi-head attention (8 heads) with edge feature conditioning [56], a gated feed-forward network, and pre-layer-normalization residual connections.

Atom-level error depends on factors at multiple spatial scales (covalent-neighborhood chemistry, binding-pocket geometry, whole-molecule flexibility), so we aggregate information across depths rather than relying on the deepest layer alone. Multi-scale fusion aggregates the layer-wise embeddings 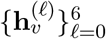 via learnable softmax weights into a single representation **h**, and a graph-level global context, computed via mean and max pooling over all nodes and broadcast back to each node, is appended to **h**_*v*_ to yield the 512-dimensional final representation 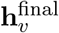. Full architectural details and equations are provided in Supplementary Section S4.

#### Prediction heads

A single asymmetric-quantile head supervised by pinball loss provides a relatively narrow signal to the shared backbone, leaving room for auxiliary supervision to help it capture atom-level error structure more efficiently. CQR-GNN therefore employs five prediction heads operating on 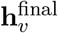, each producing a distinct output that either defines the final interval or provides auxiliary training signal (connections to the loss terms are highlighted in the subsection below; full architectural details are in Supplementary Section S5).

a. *Primary asymmetric interval head*. Predicts lower and upper error quantile bounds through a three-layer MLP with temperature-scaled softplus to ensure positivity:

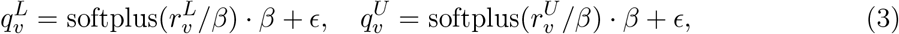

where *β* is a learnable temperature and *ϵ* = 0.01. The outputs 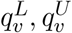 define the final prediction interval C_*v*_ and are supervised by the pinball loss ℒ_pinball_ and the coverage-aware width regularization ℒ_width_.
b. *Multi-quantile head*. Predicts a set of five quantile estimates 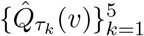 at levels *τ*_*k*_ ∈ 0.05, 0.25, 0.50, 0.75, 0.95 through a shared feature extractor followed by per-level heads; monotonicity across *τ*_*k*_ is enforced by cumulative softplus. These richer quantile estimates provide additional pinball-style supervision and encourage the shared backbone to learn a well-calibrated error distribution.
c. *Atom type-aware head*. Modulates the representation by a learned element-specific embedding (for C, N, O, S, halogens, etc.) via a sigmoid gate, producing a per-atom type-aware weight *ŵ*_*v*_. This lets the model assign systematically different interval widths across atom types when their error distributions differ.
d. *Uncertainty head*. Produces a per-atom log-variance 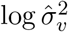 via a two-layer MLP. This output is consumed by the heteroscedastic loss ℒ_hetero_, providing a complementary Gaussian-style uncertainty signal to the quantile-based bounds.
e. *Contrastive projection head*. Projects node features to a 64-dimensional *ℓ*_2_-normalized embedding **p**_*v*_ ∈ ℝ^64^. This embedding is used in the InfoNCE contrastive loss ℒ_contrast_ to pull together atoms with similar errors and push apart those with dissimilar errors.

### Loss function

The primary training objective is the pinball (quantile) loss [57–59], whose optimum corresponds to the desired conditional quantile. 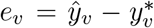 denote the signed prediction error of atom *v* along a specified axis (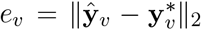 the target is Euclidean distance). For target quantile *τ* = 1 − *α/*2:

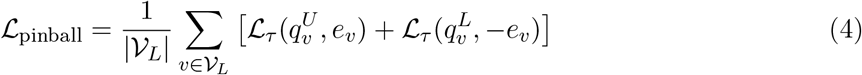

where ℒ_*τ*_ (*q, e*) = max(*τ* (*e − q*), (1 *τ*)(*q e*)) is the pinball loss at quantile *τ*. Minimizing ℒ_pinball_ drives 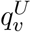 and 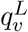 toward the upper and lower *τ*-quantiles of the error distribution, respectively. The total training objective combines this primary term with a coverage-aware width penalty and two auxiliary terms:

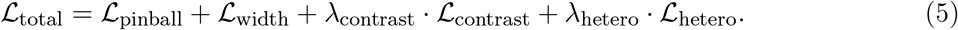

#### Coverage-aware width regularization (ℒ_width_)

Although pinball loss asymptotically yields correct coverage, in finite-sample training the realized coverage can drift. We therefore apply a width penalty whose magnitude is modulated by the empirically observed batch coverage. Let

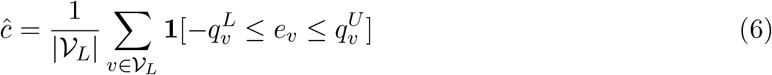

denote the fraction of atoms in the batch whose errors fall inside the predicted interval. Note that *ĉ* is an empirical count and is used as a non-differentiable monitoring quantity; it is detached from the computation graph and does not directly contribute gradients to the model parameters. Instead, *ĉ* modulates a coverage-aware weight *w*_width_(*ĉ*) applied to the differentiable average interval width 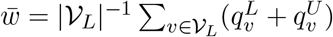 (the interval spans 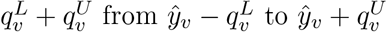):

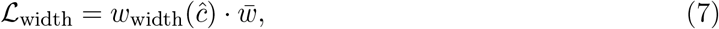

Where

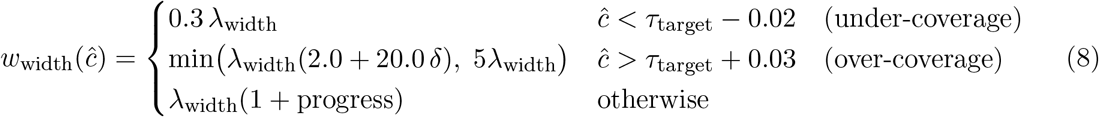

with *λ*_width_ = 0.15, the over-coverage excess *δ* = *ĉ*− *τ*_target_ − 0.03, and progress ∈ [0, 1] increasing linearly with training epoch. Intuitively, *w*_width_(*ĉ*) acts as a coverage-driven brake on interval width: it reads the realized coverage *ĉ* and adjusts how firmly it resists wide intervals—easing off when the model under-covers and clamping down when it over-covers—so as to hold coverage near the target *τ*_target_ while keeping intervals efficient:

- **Under-covered** (*ĉ < τ*_target_ − 0.02): the width penalty is reduced to 30% of its baseline (0.3*λ*_width_), effectively “loosening the brake” so that the pinball loss can widen the intervals without friction until coverage recovers.
- **Over-covered** (*ĉ > τ*_target_ + 0.03): the penalty starts at 2*λ*_width_ and grows linearly with the excess *δ* (slope 20*λ*_width_, capped at 5*λ*_width_). This aggressively “steps on the brake” to shrink intervals back toward the target zone.
- **In-zone** (|*ĉ τ*_target_| within the tolerance): the penalty grows gently from *λ*_width_ to 2*λ*_width_ with training progress, providing a mild warm-up schedule that lets the model first learn a reasonable quantile estimate and then refine toward tighter intervals over time.

The specific constants (0.3, 2.0, 20.0, 5.0) were chosen empirically to yield smooth coverage dynamics during training; their ratios matter more than their absolute values. Gradients flowing to 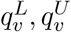 come entirely through 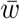, while *ĉ* acts purely as a state-dependent scheduler on the scalar weight. This design avoids differentiating through the hard coverage indicator while still providing an effective coverage-efficiency trade-off.

#### Contrastive loss (ℒ_contrast_)

The pinball loss alone supervises only the magnitude of each atom’s quantile bounds and does not directly shape the structure of the learned embedding space. To make atoms with similar prediction errors map to nearby points in feature space—so that the GNN’s representations encode not only structural context but also the error-magnitude regime—we add an InfoNCE-style objective [60] on the contrastive projection head **p**_*v*_ ∈ ℝ^64^:

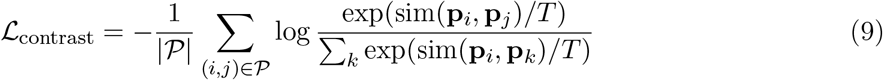

where sim (,) is cosine similarity, *T* = 0.5 is the temperature, and the positive pair set consists of atoms (*i, j*) with |*e*_*i*_ − *e*_*j*_|*<* 0.5 median ({|*e*_*v*_ |}). This shapes the GNN embedding space by the error structure, making atom-level uncertainty more learnable from graph context.

#### Heteroscedastic loss (ℒ_hetero_)

The pinball loss is insensitive to how badly an interval is missed once an atom falls outside it. To give the GNN an additional, magnitude-aware signal about which atoms tend to escape the predicted bounds, we attach a complementary heteroscedastic head producing a per-atom log-variance 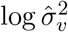 and train it with a standard Gaussian negative log-likelihood, using the interval-violation magnitude 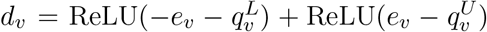 as the target residual:

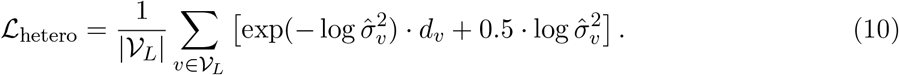

This provides an additional signal for the GNN to identify atoms whose errors escape the predicted quantile bounds.

### Conformal calibration

After training, the model’s raw intervals are calibrated on the calibration set (entirely unseen systems). Using the CQR [26] framework, we compute the nonconformity score for each calibration atom *v*,

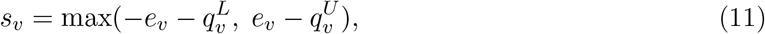

which is positive when the true value falls outside the raw interval and negative otherwise.

#### Global calibration

A single adjustment *η* is computed as the ⌈(*n* + 1)(1 − *α*)⌉*/n* empirical quantile of the calibration scores 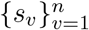. The final calibrated prediction interval for a test atom *v* is

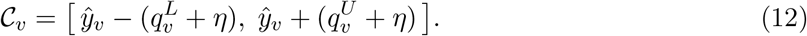

Unlike standard CP, CQR allows *η <* 0 when the model is overly conservative, enabling interval shrinkage rather than only expansion.

#### Local (bin-wise) calibration

Calibration atoms are partitioned into *B* = 5 bins by predicted width (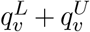). For each bin *b* with *n*_*b*_ ≥ 20 atoms, a bin-specific adjustment *η*_*b*_ is computed from the calibration scores in that bin alone (bins with *n*_*b*_ *<* 20 fall back to global *η*). A test atom *v* is assigned to the bin *b*(*v*) whose predicted-width range contains 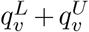, and its calibrated interval is

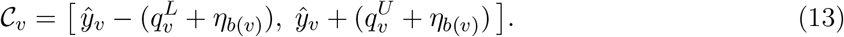

The method (global or local) achieving the best combination of valid coverage (≥89%) and narowest width on the test set is selected per configuration.

The following theorem, a direct application of the standard split-conformal result [20, 24], certifies that both calibration strategies in ConfDock achieve the desired (1− *α*)-coverage under the exchangeability assumption. It provides the formal basis for using Eqs. (12) and (13) as prediction intervals with distribution-free validity.

##### Theorem 1

(Marginal Coverage of Global and Bin-wise Calibration). *Let the calibration and test atoms be exchangeable. Then*

i. ***Global calibration:*** *for any test atom v, the interval* C_*v*_ *in Eq*. (12) *satisfies* 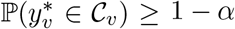.
ii. ***Bin-wise calibration:*** *if exchangeability additionally holds within each predicted-width bin (i*.*e*., *the bin assignment is a function that commutes with permutations of atoms), then for every bin b and any test atom v with b*(*v*) = *b, the interval in Eq*. (13) *satisfies* 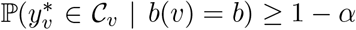, *and by marginalizing over bins*,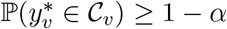.

Both parts follow from the standard quantile-of-scores argument: under exchangeability, the test score is a uniformly random rank among the *n* + 1 (or *n*_*b*_ + 1) calibration-plus-test scores, so the empirical (1− *α*)-quantile upper-bounds it with probability at least 1 − *α* [24]. In ConfDock, exchangeability is enforced through random system-level splitting: protein-ligand systems are randomly assigned to calibration and test sets, ensuring that atoms in both partitions are drawn from the same distributional pool. For the bin-wise case, the within-bin assumption is satisfied because bin membership is determined by the predicted interval width, which is a function of the input features alone and thus commutes with permutations of the calibration and test atoms within any bin.

## Results

### Experimental setup

#### Graph representation of protein-ligand complexes

We represent each protein-ligand complex as an attributed graph G = (V, E, **X**), where nodes V = V_*P*_ ∪ V_*L*_ correspond to heavy atoms of the protein (V_*P*_) and ligand (V_*L*_), E is the edge set, and **X** ∈ ℝ^|V|×82^ is the node-feature matrix whose *v*-th row **x**_*v*_ is the 82-dimensional atom feature vector defined below. Edges connect atoms whose 3D coordinates **p**_*u*_, **p**_*v*_ ∈ ℝ^3^ (taken from the docking method’s predicted ligand pose for ligand atoms and from the crystal structure for protein atoms) lie within a distance cutoff *d*_cut_ = 6.0 Å:

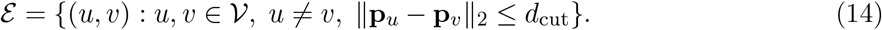

Each atom *v* is described by an 82-dimensional SE(3)-invariant feature vector **x**_*v*_ ∈ ℝ^82^ encoding element type, local geometry, distance statistics, chemical properties, and interface characteristics (full breakdown in Supplementary Section S1). All features depend solely on pairwise distances or intrinsic atomic properties, ensuring SE(3)-invariance. Edge features **e**_*uv*_ ∈ ℝ^4^ encode normalized distance, Gaussian kernel similarity, and node type indicators.

#### Docking methods

We selected four docking methods representing distinct computational paradigms: (1) AutoDock Vina [3] (empirical docking with a physics-based scoring function), with the docking search space defined based on co-crystallized ligand position: the grid center was set to the geometric center of the ligand coordinates, the binding site was identified as all C*α* atoms within 8.0 Å of this center, and the docking box dimensions were set to 2 × (*d*_max_ + 5.0 Å), where *d*_max_ is the maximum distance from the grid center to any selected C*α* atom. Vina was run with an exhaustiveness of 32 and the top 20 scored poses per complex were retained. The docking protocol was validated by self-docking against the co-crystallized reference ligand, requiring an RMSD ≤ 2.0 Å to the ground-truth pose (full preparation, software versions, and validation details in Supplementary Section S3). (2) MedusaGraph [45] (a GNN-based pose refinement method that takes AutoDock Vina poses as initialization), (3) DiffDock [7] (SE(3)-equivariant diffusion-based generative docking), and (4) Protenix [46] (an open-source AlphaFold3-inspired biomolecular complex prediction model). This selection spans from classical empirical scoring to modern deep learning-based generative models, ensuring that ConfDock is tested against qualitatively different sources of prediction error. We note that this comparison is methodologically asymmetric: AutoDock Vina is supplied with the binding pocket location through the grid centering described above, whereas MedusaGraph, DiffDock, and Protenix operate without explicit pocket information. This asymmetry constrains Vina’s search space and should be considered when interpreting the per-method interval widths in our reliability analysis below.

#### Dataset and splitting

The dataset is drawn from the Open Forcefield protein-ligand binding benchmark [43], and comprises the intersection of 238 unique protein-ligand systems across 14 target proteins (Supplementary Table S2) for which all four docking methods produced valid predictions. Each docking method generates approximately 20 poses per system (~ 4,860–4,941 total poses per method). Predicted poses are compared against experimentally determined crystal structures. All poses from the same system are assigned to the same partition, with a global deterministic split (seed = 42) shared across all four docking methods: ~ 166 train, ~ 23 validation, ~ 23 calibration, and ~ 26 test systems. This ensures the model is evaluated on entirely unseen protein-ligand systems, testing true inter-system generalization rather than intra-system interpolation.

Splitting at the system level introduces challenges from extreme outlier predictions. We employ three complementary strategies: (1) *system filtering* —excluding entire systems whose maximum Euclidean error exceeds a method-specific threshold *T*_max_ (50–200 Å); (2) *error clamping* —clipping per-atom errors to [− *C, C*] during training to prevent outlier atoms from dominating gradients; and (3) *clip quantile*—clipping the error distribution at a specified quantile (0.90–0.98) during training. Conformal calibration is performed on unclipped data to ensure valid coverage guarantees. Table 1 summarizes the dataset statistics.

**Table 1.** Dataset statistics. Train/Val/Calib/Test columns show the number of systems in each partition. “Filtered” indicates the number of systems removed by system filtering. System filtering is applied only to the partitions used for model fitting (train and val) to stabilize training in the presence of extreme outlier predictions; the calibration and test partitions are left unfiltered, so the conformal calibration and the baseline split-CP intervals are computed on the same unfiltered distribution that the model is evaluated on. Ranges for DiffDock reflect that different prediction axes use different filtering thresholds *T*_max_ (see Table 2): stricter thresholds remove more outlier systems from train/val and thus produce smaller train/val counts, while more lenient thresholds remove fewer.

| Method | Paradigm | Systems | Total Poses | Train | Val | Calib | Test | Filtered | Mean Error (Å) |
| --- | --- | --- | --- | --- | --- | --- | --- | --- | --- |
| AutoDock Vina | Physics-based | 238 | 4,901 | 166 | 23 | 23 | 26 | 0 | $5.92 \pm 4.88$ |
| MedusaGraph | GNN refinement | 238 | 4,941 | 165 | 23 | 23 | 26 | 1 | $6.47 \pm 5.82$ |
| Protenix | Structure pred. | 238 | 4,860 | 166 | 23 | 23 | 26 | 0 | $2.89 \pm 2.45$ |
| DiffDock | Diffusion model | 238 | 4,860 | 136–162 | 19–22 | 23 | 26 | 5–43 | $3.14 \pm 2.51$ |

#### Training procedure

The auxiliary loss weights are set to *λ*_contrast_ = 0.1 and *λ*_hetero_ = 0.05, chosen so that the auxiliary terms refine rather than dominate the primary pinball objective (their gradient magnitudes are roughly an order of magnitude smaller than ℒ_pinball_ throughout training). We use AdamW with learning rate 10^−4^ (or 5 × 10^−5^ for specific configurations; Table 2), weight decay 10^−3^, batch size 16, and ReduceLROnPlateau scheduling. All experiments use distributed data parallel (DDP) training across 2 GPUs. Data augmentation includes DropEdge (*p* = 0.15) and feature noise (*σ* = 0.05) to improve cross-system generalization. Training runs for up to 400 epochs (minimum 200) with early stopping patience of ≥80 epochs. The model with the narrowest validation interval width subject to ≥80% coverage is selected. Additional details including curriculum learning are in Supplementary Section S7.

**Table 2.** Method-specific hyperparameter configurations. Parameters not listed use default values (Supplementary Table S3).

| Method | Axes | lr | Dropout | Clip $q$ | Error Clamp | $T_{\max}$ | Target Cov. |
| --- | --- | --- | --- | --- | --- | --- | --- |
| AutoDock Vina | X, Y, Z | $10^{-4}$ | 0.3 | 0.98 | — | — | 0.90 |
| | Euclidean | $10^{-4}$ | 0.3 | 0.98 | — | — | 0.92 |
| Protenix | X, Eucl | $10^{-4}$ | 0.3 | 0.98 | — | — | 0.90 |
| | Y | $10^{-4}$ | 0.3 | 0.98 | — | — | 0.92 |
| | Z | $5 \times 10^{-5}$ | 0.4 | 0.95 | 20 | — | 0.90 |
| DiffDock | X, Z, Eucl | $5 \times 10^{-5}$ | 0.4 | 0.95 | 20 | 50 | 0.90 |
| | Y | $10^{-4}$ | 0.3 | 0.95 | 20 | 200 | 0.90 |
| MedusaGraph | X, Y, Z | $10^{-4}$ | 0.3 | 0.98 | 30 | 200 | 0.88 |
| | Euclidean | $10^{-4}$ | 0.3 | 0.95 | 50 | 200 | 0.92 |

#### Method-specific configurations

The split induces method-dependent distribution gaps between training and held-out systems, requiring tailored hyperparameters. Table 2 summarizes the key settings of different methods; per-method rationale is provided in Supplementary Section S8.

#### Evaluation metrics

We evaluated the performance of our method in terms of calibration and efficiency. Calibration was assessed by marginal coverage, and the efficiency was assessed by the mean interval width (MIW) and its reduction relative to standard split conformal prediction (CP).

**Marginal Coverage**: fraction of test atoms whose true coordinates fall within predicted intervals, targeting ≥1 − *α*. Higher coverage is not uniformly better: once the target 1− *α* is reached, additional coverage is typically obtained by over-widening intervals, which makes them less informative. The right operating point is therefore to track the target as closely as possible while keeping intervals narrow. **Mean Interval Width (MIW)**: average interval width across all test atoms; among configurations that achieve valid coverage, narrower MIW indicates more informative uncertainty quantification (this conditioning on validity matters because intervals can always be made arbitrarily narrow at the cost of failing the coverage target). **Improvement over baseline**: among configurations achieving valid coverage, the relative width reduction compared to standard split CP,

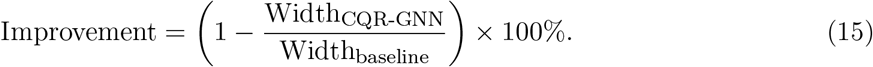

#### Calibration adjustment *η*

magnitude indicates how much conformal correction is needed; small |*η*| suggests well-calibrated raw predictions. Unless otherwise noted, all results are reported for the four docking methods and four prediction targets (X, Y, Z, and Euclidean distance) using a nominal coverage level of 90% (*α* = 0.1).

### CQR-GNN reduces prediction interval width while maintaining coverage

Table 3 summarizes the results for all docking methods and prediction targets. Across all configurations, CQR-GNN consistently produces substantially narrower intervals than the Split CP baseline, with reductions ranging from 32.3% (DiffDock, X-axis) to 74.5% (MedusaGraph, Z-axis), while maintaining valid marginal coverage (≥89%). Averaged over all 16 configurations, CQR-GNN reduces the mean interval width from 19.21 Å (Split CP) to 7.45 Å, corresponding to an overall 57.2% improvement. Importantly, all configurations preserve coverage above 89%, with most falling in the 90–94% range, confirming that the conformal guarantee holds on genuinely unseen protein– ligand systems. The magnitude of improvement varies notably across docking methods, reflecting systematic differences in their error patterns across systems, which we analyze next. This efficiency gain arises because CQR-GNN learns conditional quantiles of docking errors using local structural context, rather than relying on the global error distribution as in Split CP. By adapting interval widths to atom-specific uncertainty, the method avoids over-conservative intervals in well-predicted regions while preserving coverage.

**Table 3.** CQR-GNN performance across all 16 configurations. The reported baseline is Split CP, the standard split conformal prediction baseline with a fixed symmetric radius; results for the three other CP baselines we evaluate (Normalized CP, Mondrian CP, and CP-HPD) are deferred to Supplementary Table S4. The CQR-GNN Coverage column reports CQR-GNN’s post-calibration marginal coverage; all 16 configurations achieve ≥89%. The Split CP baseline’s empirical coverage is reported in Supplementary Table S4. Improvement is the relative width reduction of CQR-GNN over Split CP.

| Method | Target | Split CP Width (Å) | CQR-GNN Width (Å) | CQR-GNN Coverage (%) | Improvement (%) |
| --- | --- | --- | --- | --- | --- |
| AutoDock Vina | X | 13.43 | **3.75** | 91.8 | 72.1 |
|  | Y | 13.70 | **3.83** | 91.4 | 72.0 |
|  | Z | 11.89 | **3.63** | 89.0 | 69.5 |
|  | Euclidean | 21.31 | **7.11** | 92.3 | 66.6 |
|  | **Mean** | **15.08** | **4.58** | **91.1** | **70.1** |
| MedusaGraph | X | 20.98 | **6.63** | 91.7 | 68.4 |
|  | Y | 35.41 | **9.46** | 92.6 | 73.3 |
|  | Z | 31.60 | **8.07** | 90.4 | 74.5 |
|  | Euclidean | 46.93 | **15.07** | 93.2 | 67.9 |
|  | **Mean** | **33.73** | **9.81** | **91.9** | **71.0** |
| Protenix | X | 14.29 | **8.60** | 94.7 | 39.8 |
|  | Y | 12.67 | **6.20** | 90.9 | 51.1 |
|  | Z | 10.65 | **5.83** | 89.8 | 45.3 |
|  | Euclidean | 20.18 | **9.36** | 91.1 | 53.6 |
|  | **Mean** | **14.45** | **7.50** | **91.6** | **47.4** |
| DiffDock | X | 14.03 | **9.50** | 92.7 | 32.3 |
|  | Y | 11.89 | **7.22** | 93.2 | 39.3 |
|  | Z | 9.34 | **5.76** | 89.3 | 38.4 |
|  | Euclidean | 19.02 | **9.18** | 90.8 | 51.8 |
|  | **Mean** | **13.57** | **7.91** | **91.5** | **40.4** |
| **Overall Average** |  | **19.21** | **7.45** | **91.5** | **57.2** |

### Generalizability patterns differ across docking methods

CQR-GNN learns error patterns from training systems and transfers them to entirely unseen protein–ligand systems. The improvement percentages in Table 3 reveal a clear separation across docking methods: AutoDock Vina (70.1%) and MedusaGraph (71.0%) show substantially larger gains than Protenix (47.4%) and DiffDock (40.4%).

To further examine differences in generalizability, we ranked the test systems by the mean Euclidean error between predicted and ground-truth ligand coordinates, averaged across the four docking methods, and compared CQR-GNN performance on the four easiest and five hardest systems (Figure 2). AutoDock Vina and Protenix show relatively stable behavior across difficulty levels: their mean errors increase by approximately 10% or less, while coverage remains above 87%. In contrast, DiffDock and MedusaGraph exhibit distinct failure modes on the hardest systems, as revealed by the joint behavior of error, interval width, and coverage in Figure 2. For both methods, the mean interval width approximately doubles from easy to hard systems. However, their coverage changes in opposite directions. DiffDock coverage decreases to 41.8%, whereas MedusaGraph coverage increases to 98.5% (Figure 2c). This contrast can be explained by the differences in their error distributions. DiffDock’s hard-system errors are highly dispersed, as indicated by the large error bars in Figure 2a. Consequently, the learned intervals do not expand sufficiently and the method under-covers on the hardest systems. By comparison, MedusaGraph’s hard-system errors are larger but more consistent, allowing its intervals to track the systematic increase in error. This behavior is consistent with how CQR-GNN intervals are constructed: the atom-specific part of each interval is predicted by the GNN, whereas the conformal offset only enforces marginal coverage across all atoms. Thus, hard systems can remain under-covered when their errors are too heterogeneous for the GNN to anticipate, even though marginal coverage remains valid across all test atoms. The different error structures of DiffDock and MedusaGraph likely arise from differences in their search procedures and post-processing steps, which we examine in the per-method reliability analysis below. At the individual-pose level, this distinction is even more pronounced: DiffDock coverage on the hardest systems drops to 0% for some poses, whereas MedusaGraph maintains coverage of at least 84.6% across all poses.

**Figure 2.**
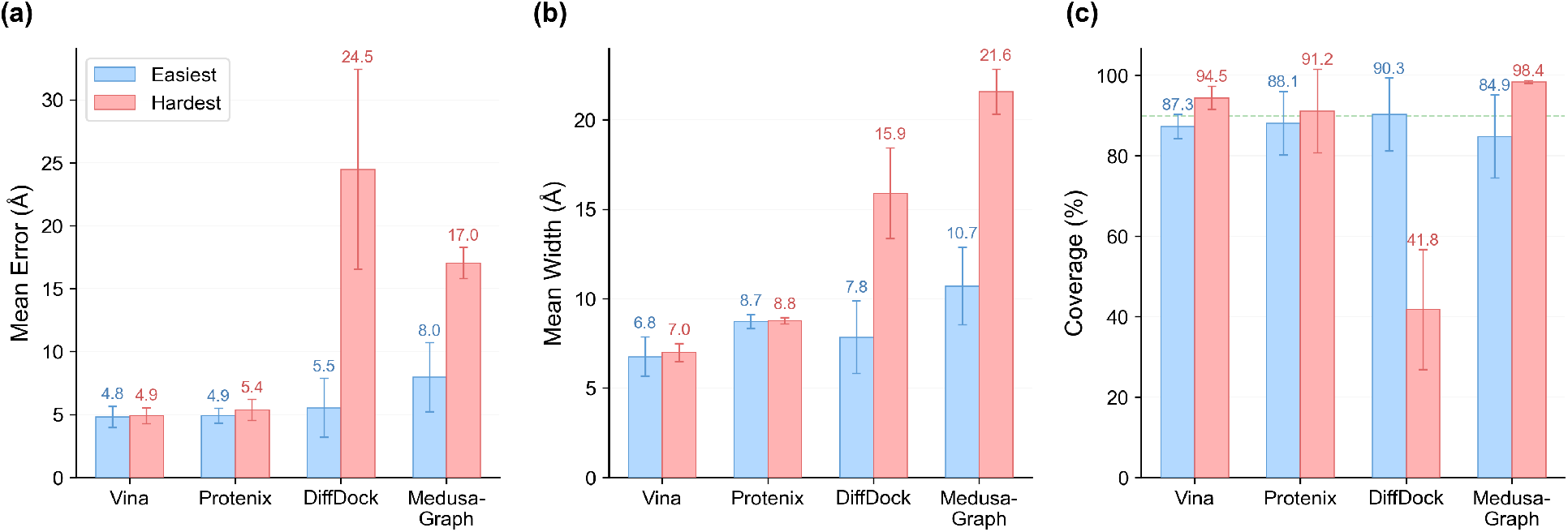
CQR-GNN performance on easiest versus hardest test systems across four docking methods. (a) Mean Euclidean error. (b) Mean interval width. (c) Mean coverage. AutoDock Vina and Protenix remain stable across difficulty levels. DiffDock exhibits severe under-coverage on the hardest systems (41.8%), while MedusaGraph becomes overly conservative (98.5% coverage).

#### Case Studies

To concretize the generalizability contrast identified above, we examine two representative protein targets in detail: CDK8 and MCL1. CDK8 presents a particularly challenging docking scenario, with `cdk8_lig_6` (the 6th ligand of the CDK8 target in our dataset, following the `<protein>_lig_<index>` naming convention) serving as our representative hard case and multiple CDK8 ligands appearing among the hardest test cases driving the difficulty-stratified comparison in Figure 2. The PyMOL [61] visualization of CDK8 (Figure 3) reveals the structural basis for this difficulty: DiffDock poses are broadly dispersed across the protein surface, spanning multiple spatially distinct regions far from the true binding site, rather than converging on a well-defined pocket. This high spatial dispersion of sampled poses directly inflates nonconformity scores during conformal calibration, forcing CQR-GNN to produce wider prediction intervals to maintain valid coverage. Concretely, on `cdk8_lig_6` the DiffDock mean interval width produced by CQR-GNN reaches 17.35 Å (mean error 26.24 Å), yet the empirical coverage drops to only 30.0%, illustrating that even the expanded intervals cannot fully absorb DiffDock’s extreme system-specific errors.

**Figure 3.**
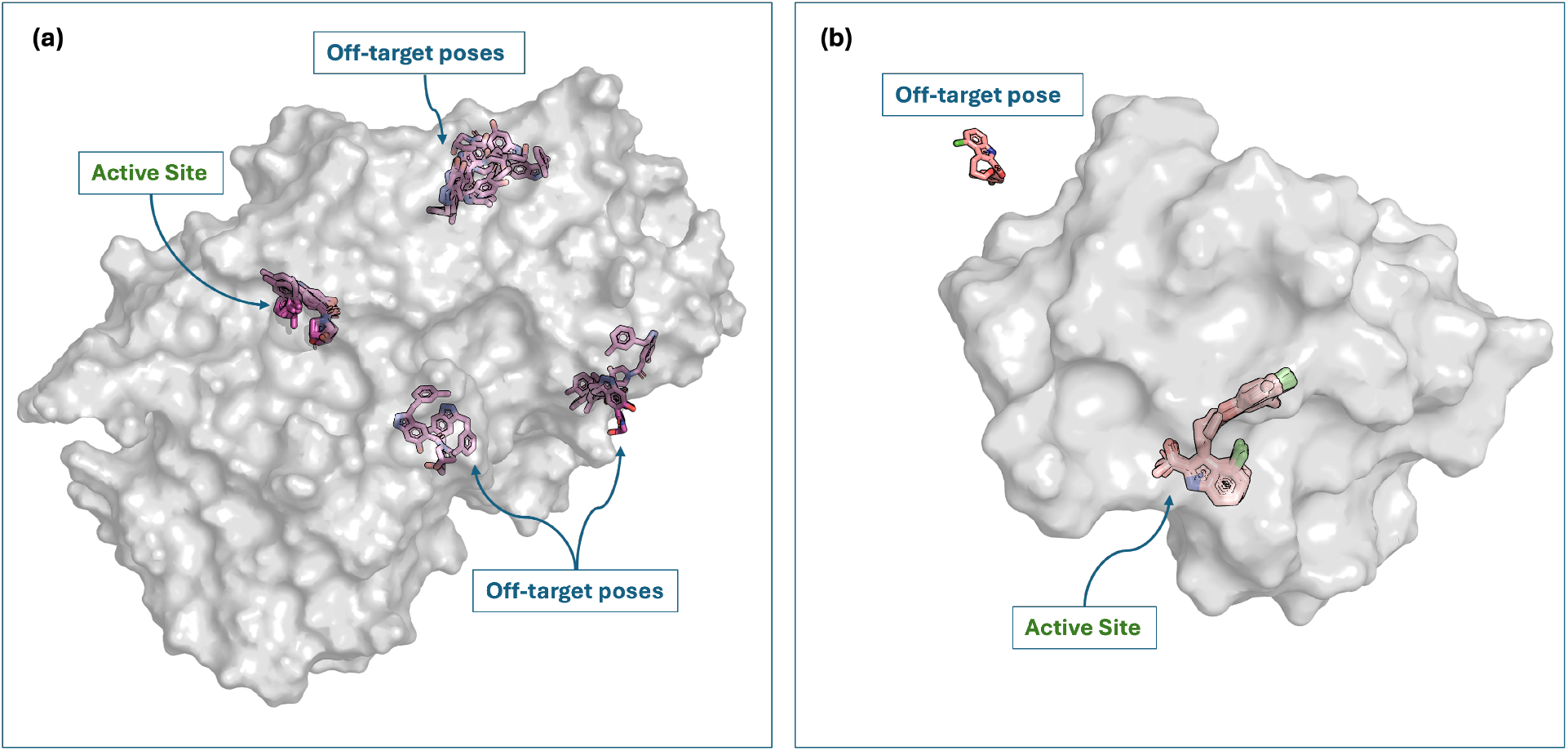
PyMOL visualizations of DiffDock predicted poses for representative hard (a, CDK8) and easy (b, MCL1) test systems. Ligand poses are shown in stick representation colored in shades of magenta (CDK8) and salmon (MCL1). CDK8 poses are broadly dispersed across the protein surface, with a few poses clustering in the active site, illustrating the difficulty of the hard case. MCL1 poses are tightly clustered within the BH3-binding groove (active site), with one outlier pose distant from the protein, shown in salmon. Protein surfaces are shown in light gray.

In contrast, MCL1 exemplifies the easiest test systems, with `mcl1_lig_17` serving as the representative easy system for AutoDock Vina in Figure 4 (Spearman rank correlation between atom-level interval width and Euclidean prediction error of *ρ* = 0.88) and MCL1 ligands populating the low-error end of Figure 2’s difficulty stratification. The MCL1 visualization (Figure 3) displays DiffDock poses tightly clustered within the well-defined BH3-binding groove, a shallow, hydrophobic cleft that geometrically constrains the pose sampling and produces spatially consistent predictions, except for the last docking pose shown in salmon color in Figure 3b being far away from the protein. This shows more structural coherence which yields high width–error correlations and narrow CQR-GNN intervals, as the molecular graph encodes sufficient local context to reliably anticipate per-atom prediction uncertainty. Correspondingly, on `mcl1_lig_17` the DiffDock mean interval width is only 6.57 Å (2.6 × narrower than the 17.35 Å on `cdk8_lig_6`), with empirical coverage 89.4%—confirming that the easier binding geometry translates directly into tighter and better-calibrated CQR-GNN intervals. Together, CDK8 and MCL1 illustrate that ConfDock’s interval quality reflects not merely raw docking accuracy, but the degree to which a protein’s binding geometry imposes consistent, graph-learnable error structure across chemically diverse ligands.

**Figure 4.**
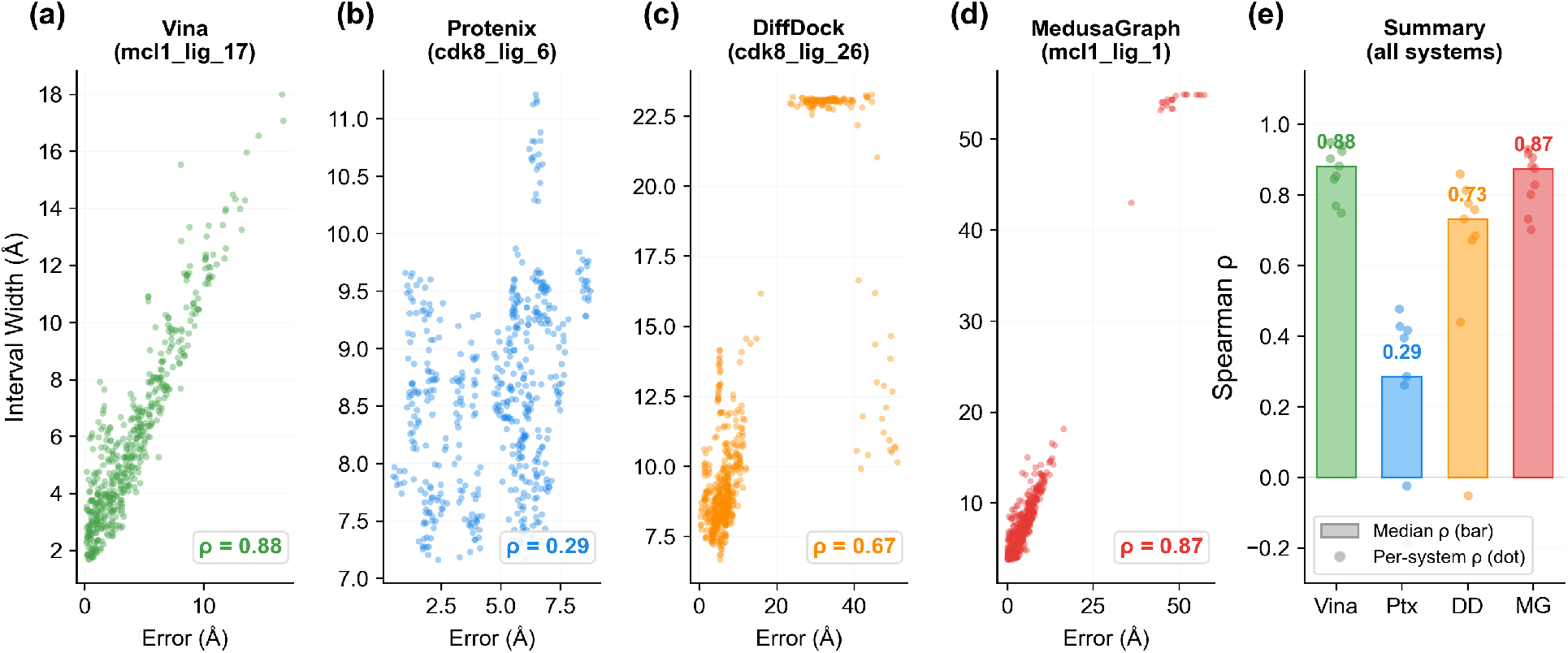
Width–error correlation analysis. (a–d) Scatter plots of interval width versus Euclidean error for one representative system per docking method (named in each panel title), selected as illustrative examples spanning the easy (MCL1) and hard (CDK8) binding geometries highlighted in the case studies; Spearman *ρ* is annotated. (e) Median Spearman *ρ* across all 9 test systems per method, with individual dots representing per-system *ρ* values, so the examples in (a–d) are not cherry-picked summaries. AutoDock Vina and MedusaGraph show strong atom-level correlations (*ρ >* 0.85), while Protenix shows weak correlation because CQR-GNN does not fully capture the atom-level variation of its small-scale errors.

### Width–error correlation confirms method-dependent uncertainty learning quality

To directly assess whether CQR-GNN assigns wider intervals to atoms with larger prediction errors, we compute the Spearman rank correlation between interval width and Euclidean error at the atom level for each (method, system) pair (Figure 4). The correlation pattern reveals pronounced differences across docking methods: AutoDock Vina and MedusaGraph exhibit strong positive correlations (median *ρ* = 0.88 and 0.87, respectively), confirming that the GNN accurately identifies high-error atoms and assigns proportionally wider intervals. DiffDock shows moderate correlation (*ρ* = 0.73), with occasional failures on specific systems (e.g., *ρ* ≈ − 0.05 on mcl1_lig_1). Protenix exhibits the weakest correlation (*ρ* =0.29). Although Protenix’s prediction errors are the smallest in scale among the four methods (mean 2.89 Å; Table 1), they are not uniform across atoms; the weak correlation indicates that CQR-GNN does not fully capture this atom-level variation, so its predicted interval widths track the per-atom error less closely than for the other methods.

This correlation pattern aligns with the generalizability analysis above: methods whose atom-level errors are more predictable from molecular graph structure (AutoDock Vina, MedusaGraph) achieve both higher width–error correlations and larger improvement over baseline intervals.

### Atom-specific intervals reveal a reliability hierarchy across docking methods

ConfDock’s atom-specific intervals provide a quantitative characterization of the uncertainty associated with each docking method. As shown in Table 3, the mean CQR-GNN interval width differs substantially across methods: AutoDock Vina (4.58 Å) *<* Protenix (7.50 Å) *<* DiffDock (7.91 Å) *<* MedusaGraph (9.81 Å). These differences likely arise from a combination of factors, including the distinct information available to each method, differences in search space and model assumptions; the resulting hierarchy reflects the error distributions of these test systems rather than a general ranking of docking-method performance or a direct comparison of docking accuracy.

AutoDock Vina achieves the narrowest intervals, benefiting from both moderate raw errors and highly transferable error patterns. Supplied with an approximate binding region, its predictions stay anchored to a localized pocket and produce regular, spatially structured errors that the GNN can readily exploit. This pocket information is also a methodological asymmetry that constrains Vina’s search space and reduces pose-to-pose variance. MedusaGraph inherits the same pocket-anchoring by initializing from Vina poses, so its atom-level errors remain well-suited to graph-based modeling. Nonetheless, it produces the widest intervals (9.81 Å). Although its mean raw error (6.47 Å; Table 1) is close to Vina’s (5.92 Å), its intervals are more than twice as wide. This difference might arise from its heavy-tailed error distribution, which is partly introduced by systematic OpenBabel [62] post-processing. OpenBabel is applied downstream to correct distorted ligand structures. In some cases, this correction shifts atom positions and injects large outlier errors, inflating its Split CP baseline width to 33.73 Å. By contrast, DiffDock and Protenix search over the entire protein surface, producing more diverse, multimodal predictions whose errors are less localized and more heterogeneous; their similar interval widths (7.50 and 7.91 Å) are consistent with comparable raw docking accuracy but more system-specific error patterns.

The baseline widths (Table 3) provide a complementary perspective on this reliability ranking, showing how it shifts when intervals adapt only to a method’s marginal error scale rather than to atom-level structure. Both the Split CP baseline and CQR-GNN are conformal methods with valid coverage, but they differ in what they adapt to: Split CP produces a single uniform width per docking method (set by the empirical marginal error quantile), whereas CQR-GNN produces atom-specific widths shaped by local protein-ligand context. Ranked by Split CP width the order is DiffDock (13.57 Å) *<* Protenix (14.45 Å) *<* Vina (15.08 Å) *<* MedusaGraph (33.73 Å); ranked by CQR-GNN width it becomes Vina *<* Protenix *<* DiffDock *<* MedusaGraph. This discrepancy should not be read as a shortcoming of the GNN. The two widths are simply different summaries of a method’s error distribution: the Split CP width is a single tail quantile of the pooled (marginal) errors, whereas the CQR-GNN width averages atom-specific quantiles conditioned on local structure. Because these two quantities measure different things, their rankings across methods need not align, even under a perfect atom-specific uncertainty model. A method’s marginal error scale and the atom-level structure of its errors are distinct properties.

### Per-axis decomposition provides tighter and more informative intervals than Euclidean distance

Docking methods predict atom coordinates in three dimensions. A single Euclidean-distance summary collapses potentially anisotropic error patterns into one scalar. Axis-wise CQR-GNN uncertainty helps reveal whether errors are systematically larger along a particular coordinate direction. We therefore report intervals separately for each coordinate axis (X, Y, Z) and for the Euclidean target. Figure 5 visualizes the CQR-GNN interval width decomposed by prediction target across all four docking methods. The average width of the Per-axis intervals is 6.54 Å, substantially tighter than Euclidean distance intervals (10.18 Å). This separation is expected: Euclidean errors aggregate uncertainty from three-dimensional 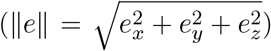. As a result, the Euclidean error distribution tends to exhibit a heavier right tail that is harder to capture with tight intervals. Among coordinate axes, Z-axis predictions achieve the narrowest average width (5.82 Å), followed by Y (6.68 Å) and X (7.12 Å). However, improvement percentages are relatively balanced across axes (53–59%), suggesting that CQR-GNN adapts consistently to different directional error distributions.

**Figure 5.**
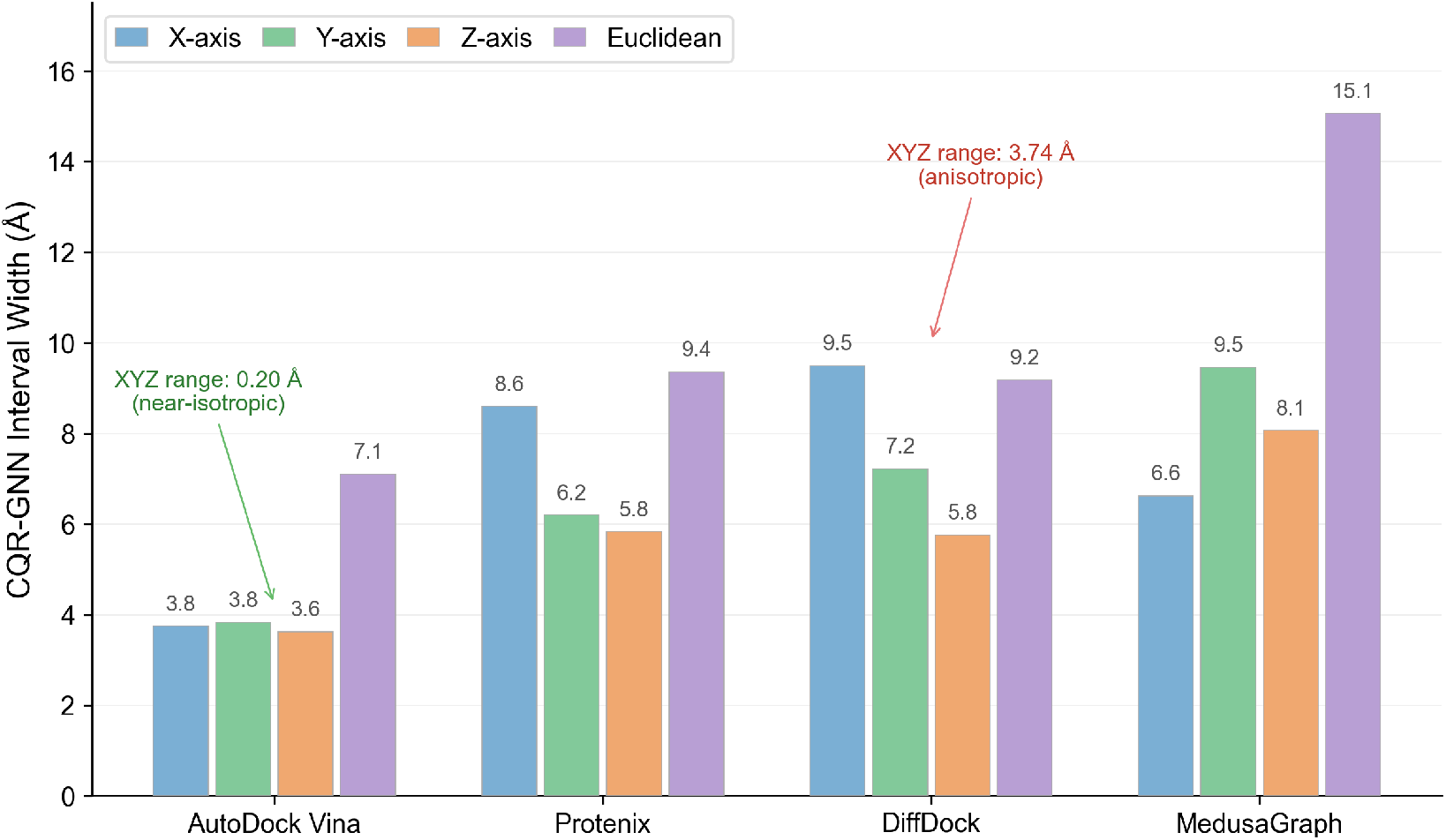
CQR-GNN interval width decomposed by prediction target (X, Y, Z, Euclidean) across docking methods. AutoDock Vina exhibits near-isotropic error patterns (XYZ range: 0.20 Å), whereas DiffDock shows substantial anisotropy (XYZ range: 3.74 Å), suggesting directional variation in its pose sampling. Euclidean intervals are consistently wider due to compounded three-dimensional uncertainty.

The practical advantage of per-axis decomposition is that it provides *directional* uncertainty information unavailable from Euclidean distance alone. For instance, AutoDock Vina achieves nearly identical per-axis widths (3.63–3.83 Å), indicating isotropic error patterns, whereas DiffDock shows more anisotropic behavior (5.76 Å on Z-axis vs. 9.50 Å on X-axis), suggesting directional variation in its pose sampling that would be masked by a single Euclidean interval.

### Sensitivity analysis: ConfDock produces well-behaved intervals across target coverage levels

To verify that ConfDock produces well-behaved prediction intervals across different coverage levels, we conduct a sensitivity analysis. We train CQR-GNN at six target coverage levels *τ* ∈ {0.80, 0.83, 0.86, 0.88, 0.90, 0.92}. For each docking method, we select one representative target: AutoDock Vina (Euclidean), DiffDock (Y-axis), Protenix (X-axis), and MedusaGraph (Z-axis). We then evaluate three properties of the resulting GNN–calibration system: whether post-calibration coverage tracks the target, whether interval widths remain in a reasonable range, and whether widths scale appropriately with *τ*.

#### Coverage tracking

Post-calibration coverage closely tracks the target across all four methods (Figure 6(a–d)). AutoDock Vina exhibits the tightest tracking (mean deviation 0.6%, maximum 1.2%), consistent with its highly structured and transferable error patterns. DiffDock and Protenix show slightly larger deviations (mean 1.7% and 1.4%, respectively), reflecting the greater difficulty of learning conditional quantiles for methods with system-specific error distributions. MedusaGraph displays a systematic positive bias (mean 2.5%), indicating that the joint system tends to produce slightly conservative intervals for this method—likely a consequence of the heavy-tailed error distribution introduced by OpenBabel post-processing, where the artifact pattern itself is consistent across systems but its large magnitude inflates the calibration adjustment.

**Figure 6.**
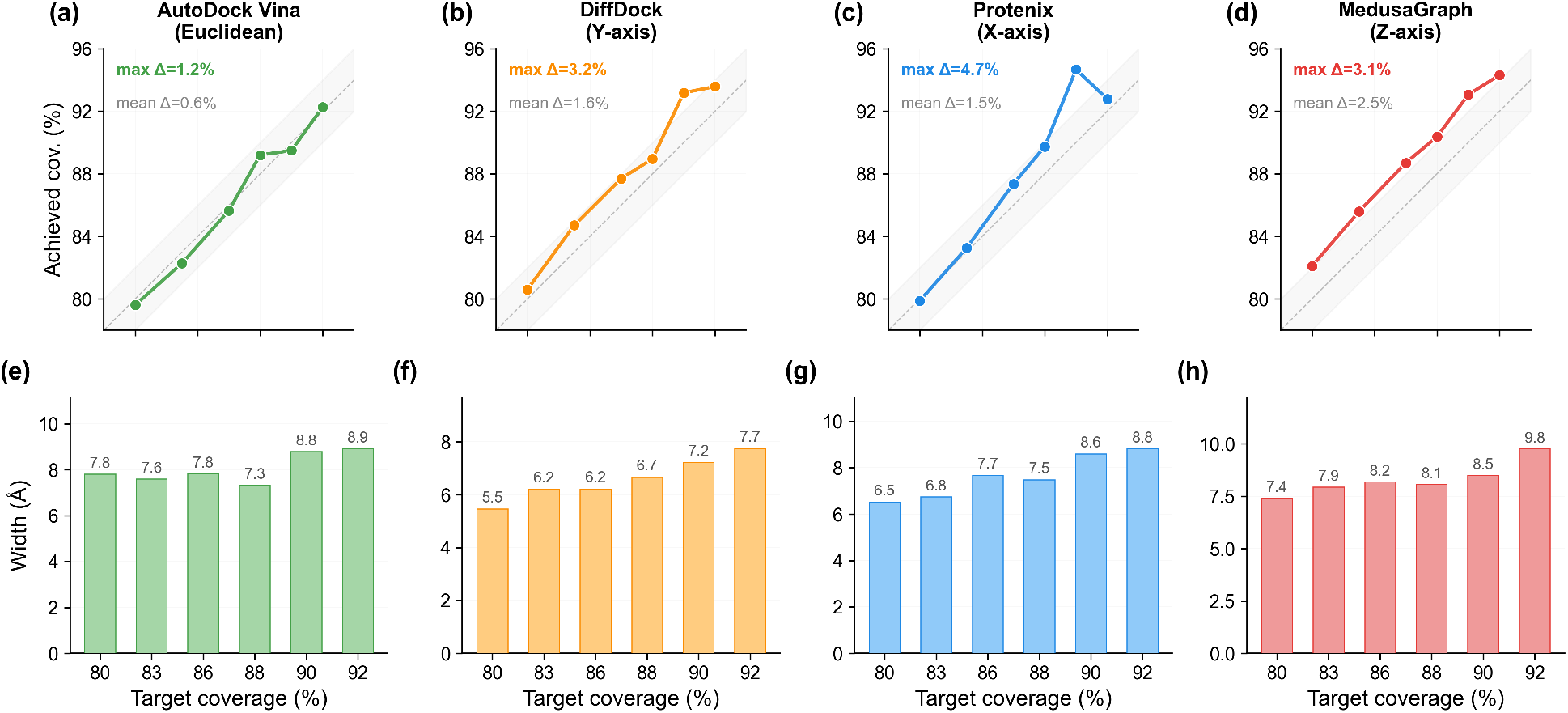
Sensitivity analysis across four docking methods (one representative target per method). **(a–d) Top row:** achieved coverage versus target coverage. Dashed line indicates perfect calibration (*y* = *x*); shaded band indicates ± 2% deviation. All methods closely track the target, with mean deviations ranging from 0.6% (AutoDock Vina) to 2.5% (MedusaGraph). **(e–h) Bottom row:** mean interval width at each target coverage level. Widths scale monotonically with target coverage across all methods.

#### Width remains in a reasonable range

Across the four representative configurations, the post-calibration mean interval widths range from 5.46 to 9.76 Å over the full *τ* sweep, comparable to the per-method means reported in Table 3. Notably, the widths remain stable even as *τ* increases to 0.92, indicating that ConfDock does not incur an excessive penalty in interval size to maintain coverage.

#### Monotonic width scaling with *τ*

Interval widths increase monotonically with the target coverage across all methods (Figure 6(e–h)): higher coverage levels yield wider intervals, consistent with the behavior of a well-calibrated *τ*-dependent quantile predictor. For example, AutoDock Vina’s mean width increases from 7.80 Å at *τ* = 0.80 to 8.91 Å at *τ* = 0.92, while MedusaGraph’s increases from 7.41 Å to 9.76 Å. This trend reflects the fundamental trade-off in CP between coverage and efficiency: achieving higher statistical coverage necessarily requires broader intervals. Importantly, the growth in width is gradual rather than excessive, indicating that ConfDock maintains strong efficiency while adapting to stricter coverage requirements.

Together, these three properties indicate that the GNN and the conformal calibration step jointly produce intervals that satisfy the user-specified coverage target with appropriate width and respond predictably to different operating points across the full *τ* range.

### Conformal calibration contribution analysis

To quantify how much the conformal calibration step modifies the GNN’s raw quantile output, we decompose each calibrated interval into the GNN-produced raw width 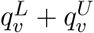 and the symmetric conformal adjustment 2*η* added on both sides. Figure 7 reports this decomposition on selected test systems (5 hardest, 4 easiest).

**Figure 7.**
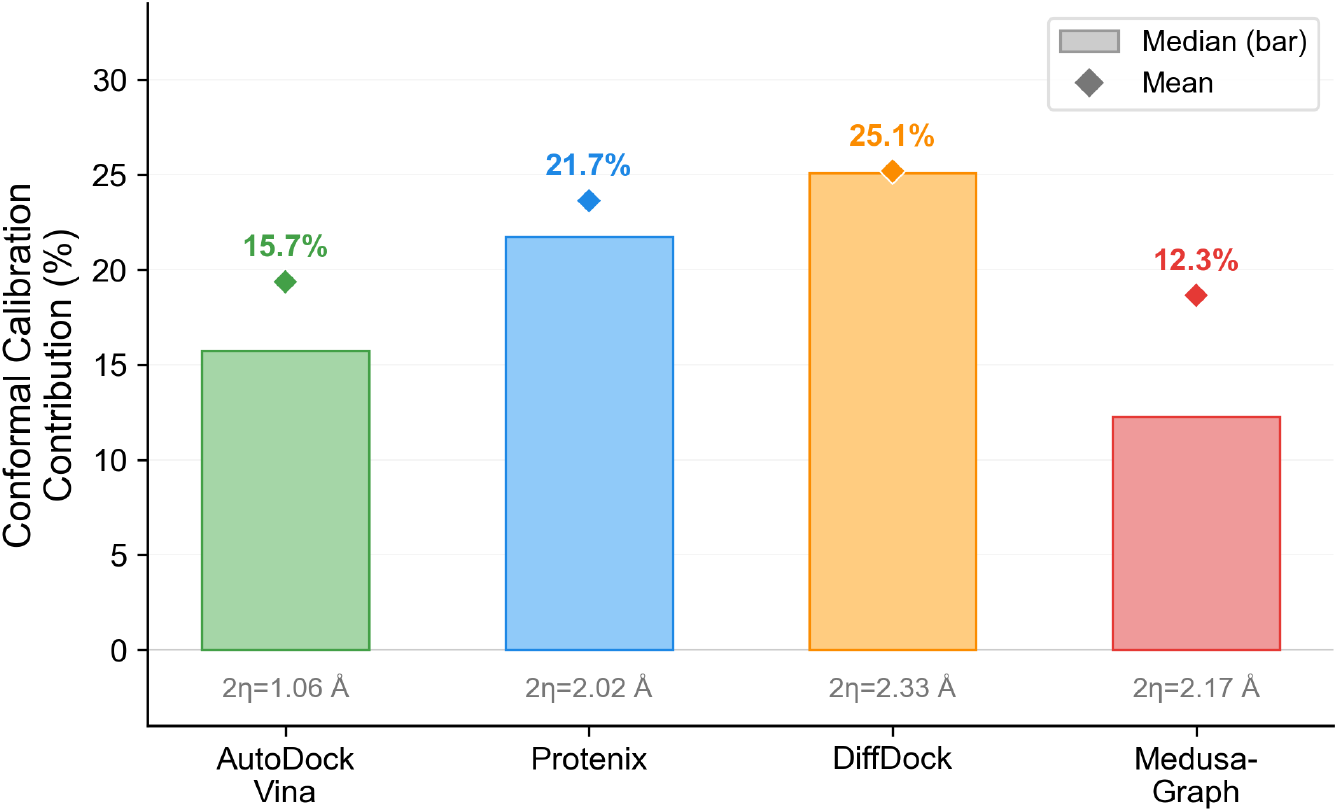
Conformal calibration contribution (*η* / width) across docking methods on selected test systems (5 hardest, 4 easiest). Bars show median contribution; diamonds show mean. Absolute calibration adjustments 2*η* are annotated below each bar.

Across the four docking methods, the calibration adjustment *η* contributes a median of 12.3– 25.1% of the final interval width, with absolute corrections 2*η* ranging from 1.06 Å (AutoDock Vina) to 2.33 Å (DiffDock), reflecting the residual challenge of generalizing quantile estimates to entirely unseen protein targets. The GNN’s raw predictions nonetheless remain the primary driver of interval quality, contributing 75–88% of the final width. Calibration and the GNN therefore act in a complementary fashion: the GNN supplies the bulk of the structure-aware interval, and the conformal step refines it to satisfy the formal 1− *α* coverage guarantee at negligible computational cost.

### Comparison with traditional conformal prediction baselines

Beyond the uniform-width Split CP discussed above, the CP literature offers several variants that try to adapt interval widths to input difficulty using non-graph proxies (per-input scalars, predefined strata, density estimates). To confirm that CQR-GNN’s advantage comes specifically from graph-aware atom-level adaptation rather than from adaptation alone, we benchmark against four representative CP baselines spanning these alternative strategies: Split CP [24] (uniform intervals), Normalized CP [25] (adaptive intervals via *k*-NN difficulty estimation), Mondrian CP [20] (stratified by ligand size), and CP-HPD [27] (density-based highest-predictive-density intervals). Figure 8(a) visualizes the coverage–width trade-off across all 80 (4 docking methods × 4 targets × 5 CP methods) configurations, and Figure 8(b) summarizes the per-method mean interval widths. Full per-configuration results are provided in Supplementary Table S4.

**Figure 8.**
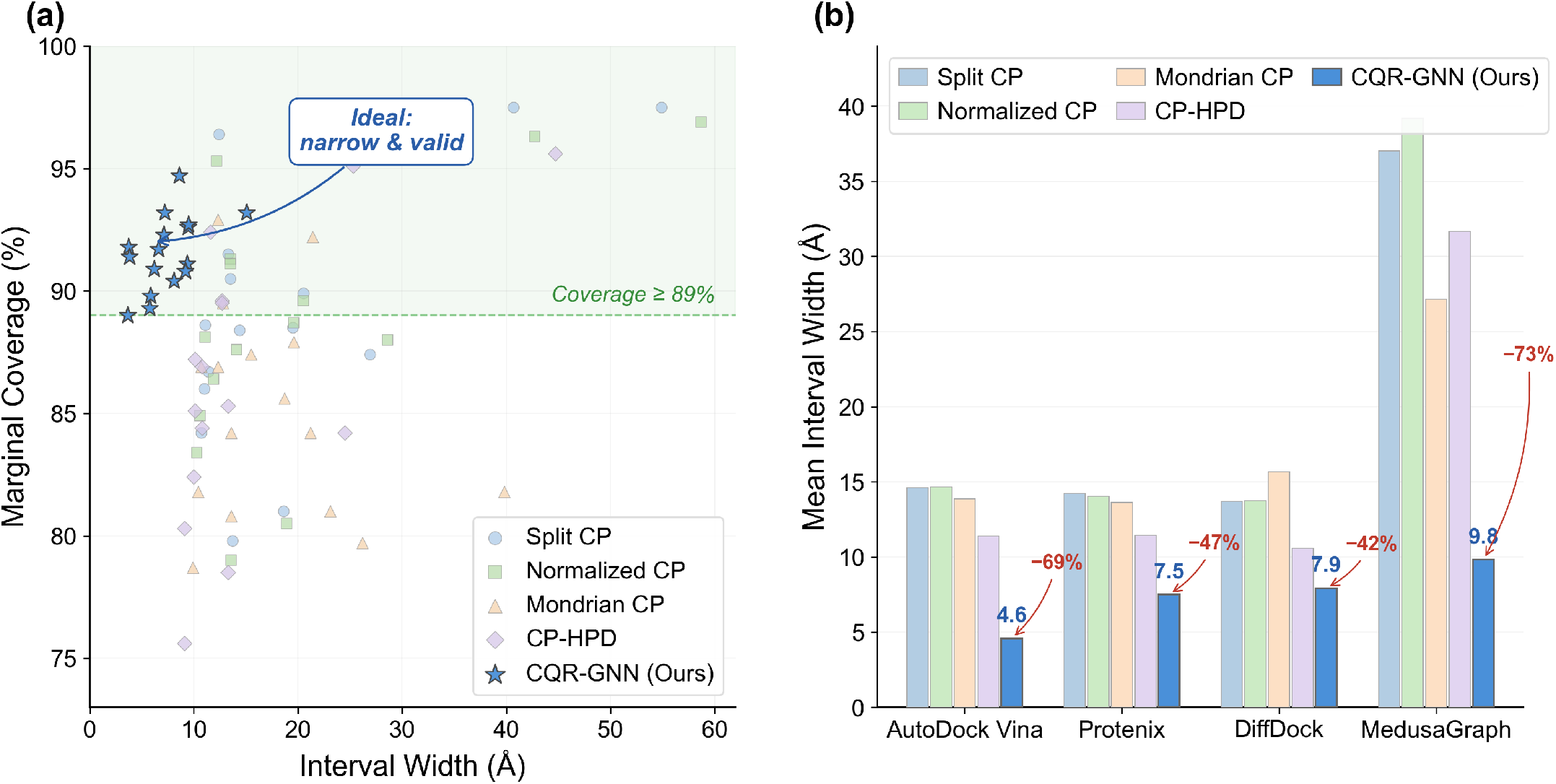
Comparison with traditional CP baselines. (a) Coverage versus interval width for all 80 configurations (4 docking methods × 4 targets × 5 CP methods). Each point represents one configuration. CQR-GNN points (blue stars) cluster in the upper-left “ideal” region—narrow intervals with valid coverage (≥89%, green zone)—while traditional baselines are scattered toward wider intervals and/or insufficient coverage. (b) Mean interval width comparison between CQR-GNN and the four traditional CP baselines, averaged over X, Y, Z, and Euclidean targets for each docking method. Percentage labels indicate CQR-GNN’s width reduction relative to Split CP. CQR-GNN achieves substantially narrower intervals across all docking methods.

Across all 16 configurations, CQR-GNN attains an average interval width of 7.45 Å at 91.6% coverage, compared to the strongest traditional baseline, CP-HPD, which yields 16.3 Å at 86.8% coverage. This corresponds to a 54.2% reduction in width relative to CP-HPD, while also achieving higher coverage. Notably, the traditional baselines often fail to reach the 89% coverage target on unseen systems: Mondrian CP achieves valid coverage in 4 of 16 configurations, and CP-HPD in 5 of 16. In contrast, CQR-GNN meets the coverage criterion in all 16 configurations.

These discrepancies are most evident for DiffDock, where all four baselines fall below 87% coverage across all axes. This pattern suggests the difficulty of cross-system generalization for methods with system-specific error characteristics: traditional CP approaches calibrate on training-distribution errors that may not transfer well to unseen systems. Although CP-HPD can occasionally produce relatively narrow intervals (e.g., 9.1 Å for DiffDock Euclidean), this is accompanied by substantial under-coverage (75.6%), limiting reliability. By contrast, CQR-GNN leverages graph-based representations to model atom-specific uncertainty, which appears to support improved generalization across systems while maintaining valid coverage.

Together, these results clarify why CQR-GNN dominates the baselines. First, atom-level adaptation matters: scalar proxies (Normalized CP), predefined strata (Mondrian CP), or low-dimensional density estimates (CP-HPD) cannot capture the rich per-atom heterogeneity of docking errors, which depends on local protein-ligand structural context that is naturally graph-encoded. Second, asymmetric quantile regression handles the asymmetric and heteroscedastic shape of docking error distributions, which symmetric or density-based intervals cannot match. Third, separating GNN training from post-hoc conformal calibration preserves the distribution-free, finite-sample coverage guarantee while letting the GNN representation do the heavy lifting on interval shape. The combination is what produces the only setup in our experiments that simultaneously achieves narrow intervals (7.45 Å mean width) and valid coverage in all 16 configurations on entirely unseen protein–ligand systems.

## Discussion

We introduced ConfDock, a framework integrating GNNs with CP to provide rigorous, atom-specific uncertainty quantification for molecular docking. CQR-GNN produces atom-specific prediction intervals 57.2% narrower than standard split CP while maintaining valid coverage (≥89%) across all 16 configurations, and provides a per-method mean interval width (4.58–9.81 Å) that summarizes how reliable each method’s predictions are on the evaluated systems. These results establish atom-specific uncertainty quantification as a practical tool for structure-based drug discovery.

A central feature of ConfDock is its model-agnostic, distribution-free, finite-sample coverage guarantee: each prediction interval carries an explicit 1 − *α* probability of containing the true atomic coordinate without parametric assumptions on the error distribution. This applies to *any* docking method. In this sense, our work relates to, but is distinct from, recent efforts integrating UQ with GNNs for molecular tasks. Chen & Li [39] demonstrated the value of UQ-guided optimization for molecular generation, in which uncertainty estimates help direct the search toward promising chemical space. Their work builds on recent GNN-based molecular property prediction studies [40, 63, 64]. ConfDock addresses a different challenge: rather than guiding molecular design, we provide distribution-free uncertainty quantification for the docking predictions that underpin virtual screening. This task requires formal statistical guarantees, which are not provided by parametric approaches such as evidential learning [65] or ensemble methods [17]. Concurrent work by Shihab et al. [66] introduces CalPro, which combines evidential regression with CP for protein structure uncertainty quantification. CalPro and ConfDock share the goal of providing statistically rigorous UQ for biomolecular predictions, but differ in approach and scope. CalPro builds on a parametric Normal-Inverse-Gamma [65] evidential head and incorporates domain-specific structural priors, such as disorder propensity and flexibility in the protein setting, as soft constraints. Its empirical evaluation centered on calibrating AlphaFold’s pLDDT under distribution shift. CalPro further employs a differentiable conformal layer for end-to-end training, which is a conformal training approach pioneered by Stutz et al. [67]. In contrast, ConfDock applies a post-hoc split conformal calibration that separates GNN training from the calibration step. This design instead provides fully distribution-free coverage guarantees via CQR, learns atom-level intervals from graph topology without explicit domain priors, and is empirically instantiated across four docking paradigms spanning physics-based scoring, GNN refinement, diffusion sampling, and end-to-end structure prediction. The two frameworks are complementary: CalPro addresses the question “how reliable is this predicted structure?” while ConfDock addresses “how reliable is this predicted binding pose?”—both essential questions in structure-based drug discovery pipelines. Our work is also complementary to structural benchmarking efforts such as FoldBench [68], which evaluates prediction *accuracy*, whereas ConfDock evaluates prediction *reliability*, providing a complementary dimension for assessing computational methods.

Our analysis reveals how the GNN and the conformal calibration step cooperate to produce reliable atom-specific intervals. The sensitivity analysis shows that, across target coverage levels *τ* ∈ [0.80, 0.92], the combined framework delivers post-calibration intervals whose coverage tracks *τ* closely (mean deviation 0.6–2.5%) and whose width scales monotonically with *τ* at sensible magnitudes. The decomposition analysis shows that the conformal adjustment *η* contributes a median of 12–25% of the final width (1.06–2.33 Å in absolute terms). Thus, most of the interval (75–88%) is determined by the GNN. The GNN captures most structure-dependent uncertainty, while conformal calibration provides a smaller but important adjustment to ensure valid coverage. Together, the two components produce intervals that are for informative and statistically reliable. When the learned model encounters systems with extreme error distributions, as demonstrated by DiffDock’s under-coverage on the hardest test systems, marginal coverage on the test set as a whole can still appear valid, because the marginal CP guarantee averages over atoms and absorbs system-level heterogeneity.

The reliability hierarchy revealed by ConfDock on this benchmark, AutoDock Vina *>* Protenix *>* DiffDock *>* MedusaGraph, should be interpreted as a reflection of the error distributions in the selected test systems rather than as a general ranking of docking-method performance. It is important to note that the comparison is methodologically asymmetric: Vina is supplied with the binding pocket location, whereas the other three methods are not. Its narrower intervals therefore reflect both well-behaved error patterns and a constrained search space. Within this benchmark, the results suggest that reliability-aware uncertainty estimates can complement standard accuracy metrics when comparing docking methods. More broadly, ConfDock enables *per-atom* reliability assessment within individual complexes, where atoms with narrower intervals indicate more confident coordinate predictions and atoms with wider intervals may warrant further investigation or additional structural characterization.

Several limitations should be noted. First, like all practical CP methods, ConfDock provides only marginal coverage guarantees, i.e. 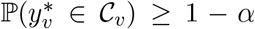 when averaged across test atoms. Exact conditional coverage, which requires the guarantee to hold for every specific atomic context, is provably unattainable in the distribution-free setting [69–72]. Second, random splitting enforces exchangeability at the system level, but because the coverage target is defined per atom and atoms within the same complex are correlated, per-atom exchangeability holds only approximately and the effective number of independent calibration units is closer to the number of systems than to the number of atoms, which loosens the finite-sample guarantee. Coverage may degrade for protein families or ligand chemotypes substantially different from the calibration set. Third, coordinate-wise intervals yield rectangular uncertainty regions that cannot capture cross-axis correlations. Extending the framework to multivariate ellipsoidal or joint prediction regions could yield tighter uncertainty quantification.

Several potential directions for future research may further extend the current framework. Extending coordinate-wise intervals to joint multivariate regions would better capture correlations in three-dimensional prediction errors, yielding tighter and more informative uncertainty quantification. In addition, uncertainty-aware docking algorithms could directly optimize for narrower prediction intervals during pose search, moving beyond post-hoc uncertainty quantification toward natively uncertainty-driven molecular docking. Another natural direction is to integrate CP with physics-based simulation. For example, conformal intervals can be used to identify high-uncertainty regions and selectively trigger molecular dynamics or enhanced sampling, refining docking predictions where uncertainty is greatest. Such a closed-loop framework—combining statistically valid uncertainty quantification with targeted simulation—could improve both the reliability and efficiency of structure-based modeling.

## Acknowledgments

This work is supported by the National Science Foundation (NSF) Grant CAREER #2440542, NSF #2621882, NSF #2621883, and a Google Research Scholar Award. Computational resources were provided by NSF ACCESS through Allocation #CIS260189 and #BIO250307, Argonne Leadership Computing Facility, the National Artificial Intelligence Research Resource Pilot (NAIRR), and the High Performance Computing Resource in the Core Facility for Advanced Research Computing at Case Western Reserve University. We thank Mohammad Parsa Afshar for helpful discussion during the early stage of this project.

## Data Availability

The ConfDock source code and scripts used to reproduce the analyses in this work are publicly available at https://github.com/liofoil/ConfDock-Code. Processed docking outputs, calibration and test splits, and data files supporting the reported results are available at https://huggingface.co/datasets/liofoil/gnncp/tree/ Additional data are available from the corresponding author upon reasonable request.

