## Supplemental Materials for "ConfDock: Atom-specific Uncertainty Quantification for Molecular Docking via Conformal Prediction"

Haochang Hao[1], Nour Elhendawy[2], Yihang Wang[2], Lu Cheng[1]
[1]University of Illinois Chicago [2]Case Western Reserve University

#### S1 Node Feature Breakdown

Each atom $v$ is described by an 82-dimensional SE(3)-invariant feature vector $\mathbf{x}_v \in \mathbb{R}^{82}$:

Table S1: Breakdown of the 82-dimensional node feature vector.

| **Category** | **Components** | **Dim** |
|---|---|---|
| Basic features | Element one-hot, residue type, protein/ligand indicators, centroid distances | 38 |
| Local geometry | Neighbor counts, anisotropy, centroid offsets at three radii | 9 |
| Distance statistics | Min/mean/std/quartile distances, radial distribution | 14 |
| Chemical properties | Electronegativity, van der Waals radius, atomic mass, H-bond potential | 5 |
| Protein-specific | Backbone, hydrophobicity, aromaticity, charge, polarity | 5 |
| Topological | Degree, clustering coefficient, 2nd-order neighbors | 3 |
| Interface | Interface flag and distance | 2 |
| Local environment | Element composition, average neighbor properties | 6 |

#### S2 Target Protein Distribution

Table S2 reports the distribution of the 238 protein–ligand systems across the 14 target proteins included in the benchmark.

Table S2: Distribution of protein-ligand systems across target proteins.

| **Target** | **Systems** | **Target** | **Systems** | **Target** | **Systems** |
|---|---|---|---|---|---|
| syk | 40 | pde2 | 18 | cdk2 | 10 |
| cdk8 | 29 | hif2a | 18 | cmet | 5 |
| mcl1 | 25 | shp2 | 17 | tnks2 | 4 |
| ptp1b | 21 | pfkfb3 | 16 | p38 | 3 |
| thrombin | 19 | tyk2 | 13 | | |

#### S3 Molecular Docking Protocol (AutoDock Vina)

**Protein and ligand preparation.** All protein structures were obtained from the Open Forcefield benchmark and preprocessed with PDBFixer to add missing atoms and generate hydrogens. Ligand molecules were converted to PDBQT format using AutoDock Tools, with polar hydrogens retained on both macromolecule and ligand.

**Grid definition.** The docking grid was defined from the spatial position of the reference co-crystallized ligand, following the procedure summarized in the main text (grid center = ligand atom centroid; binding-site C$\alpha$ atoms within 8.0 Å; box dimensions = $2 \times (d_{\max} + 5.0$ Å)).

**Docking workflow.** Each protein–ligand pair was processed in four stages:

1. **Reference pose scoring.** The experimentally observed ligand conformation was scored with the Vina empirical scoring function to establish a baseline reference energy.
2. **Local energy minimization.** The reference pose was optimized with L-BFGS to relieve steric clashes.
3. **Global search and sampling.** Monte Carlo conformational sampling was performed within the docking grid (exhaustiveness 32; energy range 3.0 kcal/mol).
4. **Post-processing.** The 20 lowest-energy poses across Monte Carlo runs were retained and ranked by predicted binding affinity.

**Validation.** Protocol accuracy was verified by self-docking each co-crystallized ligand back into its native binding pocket; success was defined as RMSD $\leq 2.0$ Å between the top-scoring predicted pose and the experimental conformation.

**Software environment.** AutoDock Vina 1.2.2; Python 3.10; BioPython 1.79 (structure parsing); NumPy 1.20+; Pandas 1.3+. A Python implementation of the workflow (`docking_pipeline.py`) is provided alongside the source code.

### S4 CQR-GNN Architecture Details

#### Input encoder

Raw node features are transformed through a two-layer MLP with layer normalization and GELU activation:

$$\mathbf{h}_v^{(0)} = W_2 \cdot \mathrm{GELU}(\mathrm{LayerNorm}(W_1 \cdot \mathrm{clamp}(\mathbf{x}_v, -2, 200) + \mathbf{b}_1)) + \mathbf{b}_2 \tag{1}$$

where $W_1 \in \mathbb{R}^{h \times 82}$, $W_2 \in \mathbb{R}^{h \times h}$, and $h = 256$ is the hidden dimension. The input clamping operation $\mathrm{clamp}(\cdot, -2, 200)$ prevents extreme feature values from outlier systems from destabilizing training.

#### Enhanced graph attention blocks

The model consists of $L = 6$ enhanced graph attention blocks, each comprising multi-head graph attention, a gated feed-forward network, and residual connections with layer normalization.

**Multi-head graph attention.** Each block applies GATv2 convolution with 8 attention heads, where each head operates on $h/8 = 32$ dimensions. Edge features are incorporated through a learned edge encoder:

$$\mathbf{e}'_{uv} = W_e^{(2)} \cdot \mathrm{GELU}(W_e^{(1)} \cdot \mathbf{e}_{uv} + \mathbf{b}_e^{(1)}) + \mathbf{b}_e^{(2)} \tag{2}$$

The attention mechanism computes:

$$\alpha_{uv}^{(k)} = \frac{\exp\left(\mathbf{a}^{(k)\top} \mathrm{LeakyReLU}\left(W_Q^{(k)} \mathbf{h}_u^{(\ell)} + W_K^{(k)} \mathbf{h}_v^{(\ell)} + \mathbf{e}'_{uv}\right)\right)}{\sum_{w \in \mathcal{N}(u)} \exp\left(\mathbf{a}^{(k)\top} \mathrm{LeakyReLU}\left(W_Q^{(k)} \mathbf{h}_u^{(\ell)} + W_K^{(k)} \mathbf{h}_w^{(\ell)} + \mathbf{e}'_{uw}\right)\right)} \tag{3}$$

with self-loops enabled.

**Gated feed-forward network.** After attention, a gated FFN processes each node independently:

$$\mathbf{f}_v = W_3 \cdot \text{GELU}(\text{Dropout}(W_4 \cdot \mathbf{h}_v + \mathbf{b}_4)) + \mathbf{b}_3 \quad \in \mathbb{R}^{2h} \tag{4}$$

$$\mathbf{g}_v = \sigma(W_g \cdot \mathbf{h}_v + \mathbf{b}_g) \quad \in \mathbb{R}^{2h} \tag{5}$$

$$\mathbf{o}_v = W_o \cdot (\mathbf{f}_v \odot \mathbf{g}_v) + \mathbf{b}_o \quad \in \mathbb{R}^{h} \tag{6}$$

where $\sigma$ denotes the sigmoid function and $\odot$ element-wise multiplication.

**Residual connections and normalization.** Both sub-layers employ pre-layer-normalization residual connections:

$$\mathbf{h}'_v = \mathbf{h}_v^{(\ell)} + \text{MultiHeadAttn}(\text{LayerNorm}(\mathbf{h}_v^{(\ell)})) \tag{7}$$

$$\mathbf{h}_v^{(\ell+1)} = \mathbf{h}'_v + \text{GatedFFN}(\text{LayerNorm}(\mathbf{h}'_v)) \tag{8}$$

##### Multi-scale fusion

Multi-scale fusion aggregates representations from all layers via learnable softmax weights:

$$\mathbf{h}_v = \sum_{\ell=0}^{L} w_\ell \cdot \mathbf{h}_v^{(\ell)}, \quad \mathbf{w} = \text{softmax}(\boldsymbol{\gamma}) \tag{9}$$

where $\boldsymbol{\gamma} \in \mathbb{R}^{L+1}$ are learnable parameters, combining local information (early layers) with global context (deep layers).

##### Global context aggregation

Graph-level context vectors are computed via mean and max pooling:

$$\mathbf{c}_\mathcal{G} = \text{MLP}\left(\left[\frac{1}{|\mathcal{V}|}\sum_{v\in\mathcal{V}} \mathbf{h}_v \;\Big\|\; \max_{v\in\mathcal{V}} \mathbf{h}_v\right]\right) \in \mathbb{R}^h \tag{10}$$

The global context is broadcast back to each node: $\mathbf{h}_v^{\text{final}} = [\mathbf{h}_v \| \mathbf{c}_\mathcal{G}] \in \mathbb{R}^{2h}$.

#### S5 Auxiliary Prediction Heads

In addition to the primary asymmetric interval head (described in the main text), CQR-GNN employs four auxiliary prediction heads operating on $\mathbf{h}_v^{\text{final}} \in \mathbb{R}^{512}$:

**Multi-quantile head.** Predicts five quantiles $\tau \in \{0.05, 0.25, 0.50, 0.75, 0.95\}$ through a shared feature extractor followed by per-quantile heads:

$$\mathbf{z}_v = \text{GELU}(\text{LayerNorm}(W_{\text{shared}} \cdot \mathbf{h}_v^{\text{final}})), \quad \hat{Q}_{\tau_k} = W_{\tau_k}^{(2)} \cdot \text{GELU}(W_{\tau_k}^{(1)} \cdot \mathbf{z}_v) \tag{11}$$

Monotonicity is enforced via cumulative softplus: $\hat{Q}_{\tau_k} = \sum_{j=1}^{k} \text{softplus}(\hat{Q}_{\tau_j}^{\text{raw}})$.

**Atom type-aware head.** Incorporates element-specific priors through learned atom-type embeddings:

$$\mathbf{t}_v = \text{Embedding}(\text{type}(v)), \quad \mathbf{g}_v = \sigma(W_t[\mathbf{h}_v^{\text{final}} \| \mathbf{t}_v]), \quad \hat{w}_v = \text{softplus}(W_a(\mathbf{g}_v \odot [\mathbf{h}_v^{\text{final}} \| \mathbf{t}_v])) \tag{12}$$

**Uncertainty head.** Produces a heteroscedastic uncertainty estimate:

$$\log \hat{\sigma}_v^2 = W_u^{(2)} \cdot \text{GELU}(W_u^{(1)} \cdot \mathbf{h}_v^{\text{final}}) \tag{13}$$

**Contrastive projection head.** Projects to a 64-dimensional $\ell_2$-normalized space:

$$\mathbf{p}_v = \frac{W_c^{(2)} \cdot \text{GELU}(W_c^{(1)} \cdot \mathbf{h}_v^{\text{final}})}{\|W_c^{(2)} \cdot \text{GELU}(W_c^{(1)} \cdot \mathbf{h}_v^{\text{final}})\|_2} \in \mathbb{R}^{64} \tag{14}$$

### S6 Auxiliary Loss Formulations

**Coverage-aware width regularization.** The empirical batch coverage $\hat{c} = |\mathcal{V}_L|^{-1} \sum_v \mathbf{1}[-q_v^L \leq e_v \leq q_v^U]$ is computed as a hard count and detached from the computation graph; it serves purely as a non-differentiable scheduling signal. The coverage-aware weight $w_{\text{width}}(\hat{c})$ applied to the differentiable average half-width $\bar{w} = |\mathcal{V}_L|^{-1} \sum_v (q_v^L + q_v^U)$ adapts based on coverage state:

$$w_{\text{width}}(\hat{c}) = \begin{cases} 0.3\,\lambda_{\text{width}} & \hat{c} < \tau_{\text{target}} - 0.02 \quad \text{(under-coverage)} \\ \min\left(\lambda_{\text{width}}(2.0 + 20.0\delta),\ 5\lambda_{\text{width}}\right) & \hat{c} > \tau_{\text{target}} + 0.03 \quad \text{(over-coverage by } \delta) \\ \lambda_{\text{width}}(1 + \text{progress}) & \text{otherwise (in-zone)} \end{cases} \tag{15}$$

where $\lambda_{\text{width}} = 0.15$ and $\text{progress} = \min(\text{epoch}/\text{max\_epochs}, 1.0)$. The full regularizer is $\mathcal{L}_{\text{width}} = w_{\text{width}}(\hat{c}) \cdot \bar{w}$. Gradients flow only through $\bar{w}$ to the quantile bounds; $\hat{c}$ contributes no gradient but modulates the scalar weight batch-by-batch.

**Contrastive loss.** InfoNCE-style loss encouraging atoms with similar errors to have similar representations:

$$\mathcal{L}_{\text{contrast}} = -\frac{1}{|\mathcal{P}|} \sum_{(i,j)\in\mathcal{P}} \log \frac{\exp(\text{sim}(\mathbf{p}_i, \mathbf{p}_j)/T)}{\sum_k \exp(\text{sim}(\mathbf{p}_i, \mathbf{p}_k)/T)} \tag{16}$$

where $T = 0.5$ and positive pairs $\mathcal{P}$ satisfy $|e_i - e_j| < 0.5 \cdot \text{median}(\{|e_v|\})$.

**Heteroscedastic loss.**

$$\mathcal{L}_{\text{hetero}} = \frac{1}{|\mathcal{V}_L|} \sum_{v\in\mathcal{V}_L} \left[\exp(-\log \hat{\sigma}_v^2) \cdot d_v + 0.5 \cdot \log \hat{\sigma}_v^2\right] \tag{17}$$

where $d_v = \text{ReLU}(-e_v - q_v^L) + \text{ReLU}(e_v - q_v^U)$ is the interval violation magnitude.

### S7 Training Details

**Curriculum learning.** Early epochs focus on easier samples, gradually transitioning to uniform weighting:

$$w_v^{\text{curr}} = \exp\left(-2 \cdot (1 - \text{progress}) \cdot r_v\right) \tag{18}$$

where $\text{progress} = \min(\text{epoch}/T_{\text{warm}}, 1.0)$ and $r_v \in [0, 1]$ is the normalized rank of atom $v$ by error magnitude.

**Model selection criterion.**

$$\text{score} = \bar{w}_{\text{val}} \cdot \begin{cases} 1 + 5 \cdot (0.80 - \hat{c}_{\text{val}}) & \text{if } \hat{c}_{\text{val}} < 0.80 \\ 1.0 & \text{otherwise} \end{cases} \tag{19}$$

The model with the lowest score (narrowest intervals subject to reasonable coverage) is selected.

### S8 Method-Specific Configuration Rationale

**AutoDock Vina.** Cleanest dataset (max Euclidean error = 23.7 Å). No system filtering or error clamping needed. Euclidean axis uses elevated target coverage (0.92) for valid post-calibration coverage.

**Protenix.** Similarly clean (max Euclidean error = 18.8 Å). Z-axis benefits from lower learning rate ($5 \times 10^{-5}$) and higher dropout (0.4) to handle a wider train-test distribution gap, with error clamping at 20 Å.

**DiffDock.** Noisiest dataset with extreme outliers (max Euclidean error up to $9.4 \times 10^7$ Å). System filtering ($T_{\max} = 50$ Å) removes 43 pathological systems for X/Z/Euclidean; Y-axis uses $T_{\max} = 200$ Å (5 systems removed). Error clamping at 20 Å, lower learning rate, and higher dropout are essential.

**MedusaGraph.** Moderate noise (max Euclidean error = 269 Å, median = 9.6 Å). One system removed by $T_{\max} = 200$ Å. Error clamping (30 Å per-axis, 50 Å Euclidean) and reduced target coverage (0.88 per-axis) balance tightness with post-calibration validity.

### S9 Complete Hyperparameter Summary

Table S3 consolidates the default values and tuning ranges for all hyperparameters used by Conf-Dock; method-specific overrides are listed in Table 2 of the main text.

### S10 Diagnostic Coverage Analysis by Element Type

While conformal prediction provides only marginal coverage guarantees, it is important to verify that adaptive methods do not introduce systematic biases for specific atom types. Figure S1 shows the coverage breakdown by element type across four docking methods on selected test systems (5 hardest, 4 easiest).

AutoDock Vina and MedusaGraph maintain coverage above 80% for all major element types (C, N, O, S), with no evidence of systematic element-specific bias. Protenix shows anomalously low coverage for sulfur atoms (45.0%), though the sample size is small ($n = 40$) and sulfur atoms in ligand functional groups (e.g., sulfonamides) tend to occupy solvent-exposed positions with inherently higher prediction uncertainty. DiffDock exhibits uniformly low coverage across all elements (59–65%), which reflects its catastrophic failure on the hardest cdk8 systems rather than element-specific bias—consistent with the difficulty-stratified analysis in the main text.

Table S3: Complete hyperparameter summary.

| Category | Parameter | Default / Range | Description |
|---|---|---|---|
| Architecture | Hidden dimension | 256 | GNN hidden dimension |
| | # GAT layers | 6 | Number of attention blocks |
| | # Attention heads | 8 | Heads per block |
| Optimization | Learning rate | $10^{-4}$ ($5 \times 10^{-5}$–$10^{-4}$) | Initial learning rate |
| | Weight decay | $10^{-3}$ | AdamW weight decay |
| | Batch size | 16 | Graphs per batch |
| | Dropout | 0.3 (0.3–0.4) | Dropout rate |
| | Max / Min epochs | 400 / 200 | Training duration |
| Loss weights | $\lambda_{\text{width}}$ | 0.15 | Width regularization |
| | $\lambda_{\text{coverage}}$ | 0.5 | Coverage loss weight |
| | $\lambda_{\text{contrastive}}$ | 0.1 | Contrastive loss weight |
| | $\lambda_{\text{hetero}}$ | 0.05 | Heteroscedastic loss weight |
| Preprocessing | Target coverage | 0.90 (0.88–0.93) | Training target $1 - \alpha$ |
| | Clip quantile | 0.98 (0.90–0.98) | Error clip percentile |
| | Error clamp | — (15–100) | Hard error clamp (Å) |
| | System threshold | — (50–200) | Filtering threshold (Å) |
| Augmentation | DropEdge rate | 0.15 | Edge removal probability |
| | Feature noise std | 0.05 (0.05–0.10) | Gaussian noise std |
| | Contrastive temp. | 0.5 | InfoNCE temperature |
| Early stopping | Patience | 80 (80–100) | Epochs without improvement |

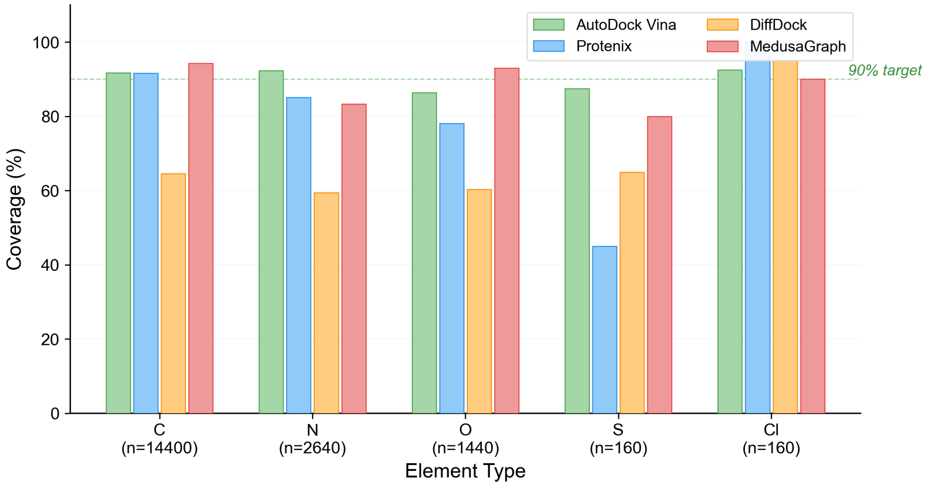


Figure S1: Coverage by element type across docking methods on selected test systems. Dashed line indicates the 90% target. DiffDock's uniformly low coverage reflects whole-system failures rather than element-specific bias.

#### S11 Detailed Baseline Comparison

Table S4 reports the per-configuration interval width and empirical coverage for CQR-GNN and the four traditional CP baselines (Split CP, Normalized CP, Mondrian CP, CP-HPD) summarized in the main text.

Table S4: Interval width (Å) and coverage (%) comparison between CQR-GNN and traditional CP baselines across all 16 configurations. Bold indicates the narrowest width per row among methods achieving ≥89% coverage. †Coverage below 89%.

| **Method** | **Target** | **Split CP**<br>W / Cov | **Norm. CP**<br>W / Cov | **Mondrian**<br>W / Cov | **CP-HPD**<br>W / Cov | **CQR-GNN**<br>W / Cov |
|---|---|---|---|---|---|---|
| AutoDock Vina | X | 13.43 / 91.5 | 13.47 / 91.3 | 12.76 / 89.5 | 12.65 / 89.6 | **3.75** / 91.8 |
| | Y | 13.70 / 90.5 | 13.45 / 91.1 | 12.33† / 86.9 | 12.65 / 89.5 | **3.83** / 91.4 |
| | Z | 11.89† / 88.6 | 11.11† / 88.1 | 10.71† / 86.9 | 10.12† / 85.1 | **3.63** / 89.0 |
| | Eucl | 21.31 / 89.9 | 20.48 / 89.6 | 19.60† / 87.9 | 10.12† / 87.2 | **7.11** / 92.3 |
| MedusaGraph | X | 20.98 / 96.7 | 26.68 / 96.4 | 21.45 / 92.2 | 25.31 / 95.1 | **6.63** / 91.7 |
| | Y | 35.41 / 97.5 | 42.69 / 96.3 | 26.16† / 79.7 | 44.71 / 95.6 | **9.46** / 92.6 |
| | Z | 31.60† / 87.4 | 28.57† / 88.0 | 21.24† / 84.2 | 24.47† / 84.2 | **8.07** / 90.4 |
| | Eucl | 46.93 / 97.5 | 58.74 / 96.9 | 39.84† / 81.8 | 32.06 / 96.8 | **15.07** / 93.2 |
| Protenix | X | 14.29† / 88.4 | 14.11† / 87.6 | 13.57† / 84.2 | 13.29† / 85.3 | **8.60** / 94.7 |
| | Y | 12.67† / 84.2 | 10.27† / 83.4 | 9.90† / 78.7 | 9.97† / 82.4 | **6.20** / 90.9 |
| | Z | 10.65 / 96.4 | 12.18 / 95.3 | 12.30 / 92.9 | 11.63 / 92.4 | **5.83** / 89.8 |
| | Eucl | 20.18† / 88.5 | 19.61† / 88.7 | 18.67† / 85.6 | 10.80† / 86.9 | **9.36** / 91.1 |
| DiffDock | X | 14.03† / 79.8 | 13.64† / 79.0 | 13.61† / 80.8 | 13.29† / 78.5 | **9.50** / 92.7 |
| | Y | 11.89† / 86.7 | 11.87† / 86.4 | 10.36† / 81.8 | 10.80† / 84.4 | **7.22** / 93.2 |
| | Z | 9.34† / 86.0 | 10.64† / 84.9 | 15.53† / 87.4 | 9.14† / 80.3 | **5.76** / 89.3 |
| | Eucl | 19.02† / 81.0 | 18.91† / 80.5 | 23.11† / 81.0 | 9.14† / 75.6 | **9.18** / 90.8 |
| **Average** | | 19.21 / 89.4 | 20.40 / 89.0 | 17.57† / 85.1 | 16.26† / 86.8 | **7.45** / 91.6 |

#### S12 Sensitivity Analysis: Detailed Results

Table S5 provides the complete sensitivity analysis data for CQR-GNN at six target coverage levels across four docking methods (one representative prediction target per method).

Table S5: Sensitivity analysis: CQR-GNN achieved coverage and mean interval width at varying target coverage levels. One representative prediction target is shown per docking method. Δ: deviation from target coverage.

| **Method (Target)** | $\tau$ | **Achieved Cov. (%)** | **Δ (%)** | **Width (Å)** | **Epochs** |
|---|---|---|---|---|---|
| AutoDock Vina (Euclidean) | 0.80 | 79.6 | −0.4 | 7.80 | 200 |
| | 0.83 | 82.3 | −0.7 | 7.59 | 200 |
| | 0.86 | 85.7 | −0.4 | 7.82 | 200 |
| | 0.88 | 89.2 | +1.2 | 7.33 | 260 |
| | 0.90 | 89.5 | −0.5 | 8.79 | 200 |
| | 0.92 | 92.3 | +0.3 | 8.91 | 330 |
| DiffDock (Y-axis) | 0.80 | 80.6 | +0.6 | 5.46 | 317 |
| | 0.83 | 84.7 | +1.7 | 6.21 | 276 |
| | 0.86 | 87.7 | +1.7 | 6.22 | 324 |
| | 0.88 | 89.0 | +1.0 | 6.66 | 322 |
| | 0.90 | 93.2 | +3.2 | 7.22 | 347 |
| | 0.92 | 93.6 | +1.6 | 7.74 | 400 |
| Protenix (X-axis) | 0.80 | 79.9 | −0.1 | 6.53 | 250 |
| | 0.83 | 83.3 | +0.3 | 6.75 | 294 |
| | 0.86 | 87.4 | +1.4 | 7.68 | 253 |
| | 0.88 | 89.7 | +1.7 | 7.49 | 293 |
| | 0.90 | 94.7 | +4.7 | 8.60 | 251 |
| | 0.92 | 92.8 | +0.8 | 8.82 | 346 |
| MedusaGraph (Z-axis) | 0.80 | 82.1 | +2.1 | 7.41 | 367 |
| | 0.83 | 85.6 | +2.6 | 7.93 | 313 |
| | 0.86 | 88.7 | +2.7 | 8.19 | 336 |
| | 0.88 | 90.4 | +2.4 | 8.07 | 400 |
| | 0.90 | 93.1 | +3.1 | 8.49 | 400 |
| | 0.92 | 94.3 | +2.3 | 9.76 | 373 |
